# Neuronal Toll-like Receptor-4 regulation of Matrix Metalloproteinase-9 Activity Mediates Dentate Circuit Dysfunction after Traumatic Brain Injury

**DOI:** 10.64898/2026.01.09.698505

**Authors:** Deepak Subramanian, Erick Contreras, Laura Dovek, Razieh Jaberi, Emmanuel Green, Ysabelle K Lao, Iryna M Ethell, Vijayalakshmi Santhakumar

## Abstract

Neuroinflammatory pathways activated by traumatic brain injury (TBI) are critical mediators of long-term neurological dysfunction and represent promising therapeutic targets. Toll-like receptor 4 (TLR4), an innate immune receptor, was previously shown to contribute to increased seizure susceptibility and cognitive deficits in rats after lateral fluid percussion injury (FPI). However, the cellular and molecular mechanisms underlying TLR4-mediated circuit dysfunction early after brain injury are not fully understood. In this study, we define a cell- and circuit-specific neuroimmune-enzyme effector signaling axis that mediates early post-TBI circuit dysfunction in the hippocampal Dentate Gyrus (DG). Using *ex vivo* electrophysiology in rat and mouse models one-week after brain injury, we demonstrate that neuronal TLR4 signaling regulates both excitatory and inhibitory synaptic inputs to dentate granule cells (DGC). Collectively, pharmacological inhibition of TLR4 in rats and cell-type-specific deletion of TLR4 in mice show that neuronal TLR4 mediates injury-driven increase in DGC excitatory input frequency and relies on downstream activation of Matrix Metalloproteinase-9 (MMP-9). In contrast, TLR4 signaling contributed to decrease in inhibitory current frequency after injury, but independent of MMP-9, revealing a mechanistic divergence. Systemic inhibition of either TLR4 signaling or MMP-9 activity in rats within 24 hours after injury reduced network hyperexcitability and improved long-term potentiation (LTP) in the DG measured *in vivo* one-week after injury. Either TLR4 or MMP-9 inhibition early after injury effectively attenuated spatial memory deficits in a Barnes maze task one-month post-injury. Paradoxically, in sham controls, inhibition of TLR4 increased frequency of both excitatory and inhibitory inputs to DGCs and augmented network excitability, without altering MMP-9 levels, identifying context-dependent roles for TLR4 signaling. Together, these results identify a novel TLR4-MMP-9 axis as a key driver of early post-TBI dentate gyrus circuit dysfunction and behavioral deficits.

## INTRODUCTION

Traumatic brain injury (TBI) initiates a robust and complex inflammatory cascade that exacerbates secondary structural and functional damage [1–4]. While cellular damage is widespread and diffuse, the hippocampus is highly susceptible to post-traumatic pathology. Specifically, the hippocampal dentate gyrus (DG) undergoes extensive loss of hilar neurons and circuit reorganization that critically alter hippocampal network physiology and contributes to the development of cognitive impairments and posttraumatic epilepsy [5–11]. In this context, we previously identified a pivotal role for the neuronally expressed innate immune receptor, Toll-like receptor 4 (TLR4), in enhancing neuronal excitability and driving epileptogenesis after injury [12–14]. Yet, the downstream mechanisms by which specific immune receptors influence circuit physiology remain poorly understood.

Recent evidence indicates that canonical TLR4 activators, such as lipopolysaccharide (LPS) and high-mobility group box 1 (HMGB1), can upregulate Matrix Metalloproteinase-9 (MMP-9) activity in vitro [15,16]. MMP-9, a zinc-dependent endopeptidase, is a critical regulator of synaptic communication and plasticity through the activity-dependent degradation of the extracellular matrix (ECM)[17–20]. Importantly, MMP-9 levels are elevated early in both animal models and clinical populations after brain injury [21–24], driving pathological remodeling of synapses and circuits that promote epileptogenesis [25–29]. Despite a strong temporal correlation between MMP-9 elevation and transient increases in TLR4 signaling after injury [12,30], a causal association between these two pathways has not been established. Based on the temporal correlation and shared upstream activators, we hypothesized that mechanistic interactions between these two pathways contribute to early circuit dysfunction after brain injury.

Here, we provide evidence for a novel mechanism involving TLR4-mediated upregulation of MMP-9 activity following brain injury that drives maladaptive changes in synaptic physiology, network excitability, and plasticity. Our findings reveal that the injury-induced increase of MMP-9 in the dentate gyrus is specifically mediated by neuronal TLR4 signaling. While TLR4-regulated MMP-9 activity was critical in enhancing excitatory synaptic inputs, inhibitory input dysregulation was mediated by TLR4 signaling independent of MMP-9.

Furthermore, we demonstrate that this TLR4 and MMP-9 dependent pathway enhances network excitability and impairs synaptic plasticity early after injury. Importantly, we show that pharmacological inhibition of either TLR4 or MMP-9 effectively mitigates these early pathological changes, highlighting the therapeutic potential of targeting this neuro-immune signaling pathway to preserve circuit function after TBI.

## METHODS

All experiments were performed in accordance with IACUC protocols approved by the University of California-Riverside, CA and in keeping with the ARRIVE guidelines. Male and female Wistar rats (P23-P24) and mice (TLR4^f/f^:CaMKIIa-CreERT2) were housed with littermates in a 12-hour light/dark cycle with access to water and food ad libidum. All mice were bred in-house. Animals were randomly assigned to either sham or FPI groups and treatment conditions.

### Fluid Percussion Injury (FPI) and treatments

Juvenile male and female Wistar rats (24 days old) were subjected to moderate (1.8-2.0 atm) lateral FPI or sham injury using standard methods [12–14]. Briefly, a 2.7 mm hole was trephined on the left side of the skull 3 mm caudal to bregma and 3.5 mm lateral from the sagittal suture. A Luer-Lok syringe hub with a 3 mm inner diameter was placed over the exposed dura and bonded to the skull with cyanoacrylate adhesive. The animals were returned to their home cage following recovery. One day later, animals were briefly anesthetized with isoflurane and attached to a fluid percussion injury device (Virginia Commonwealth University, Richmond, VA, Model 01-B) directly at the metal nozzle. A pendulum was dropped to deliver a brief (20ms) 1.9–2.0 atm impact on the dura. For sham injury, the animals underwent surgical implantation of the hub and were anesthetized and attached to the FPI device, but the pendulum was not dropped. Only rats with >10 seconds apnea after injury were included in the injury group, and only the hemisphere ipsilateral to the injury was examined.

To achieve neuron-specific deletion of TLR4, TLR4-floxed (TLR4^f/f^)mice were crossed with mice expressing inducible Cre recombinase under the control of the CamKIIa promoter (CaMKIIa-CreERT2). Cre-positive (Cre^+ve^) mice with TLR4 deletion and Cre-negative (Cre^-ve^) littermate were used in experiments. TLR4^f/f^:CaMKIIa-CreERT2 mice (8–10-week-old male and female mice) were induced with tamoxifen and subjected to lateral FPI (1.5 atm) or sham injury [31,32]. Tamoxifen (75mg/kg, i.p.) was prepared in 10% ethanol and 90% corn oil by sonicating at 37°C for 90 minutes. Mice received 3 doses of tamoxifen, 24 hours before injury, 30 minutes after injury and 24 hours after injury. TLR4 deletion in glutamatergic neurons in dentate gyrus was confirmed through immunohistochemistry (see immunohistochemistry methods below) and RNAscopy. Animals were treated with vehicle (DMSO), TLR4 antagonist (TAK-242, 0.5mg/kg, i.p., Invivogen, San Diego, CA) or MMP-9 antagonist (SB-3CT, 50mg/kg, i.p., SelleckChem, Houston, TX) at 30 minutes, 8 hours, and 24 hours after injury. The dosing strategy for TAK-242 was based on the approach previously adopted to identify functional outcomes [13,14,30,33]. Although SB3-CT has a shorter half life than TAK-242 [34], prior studies have adopted 30 minutes - 24 hour dosing strategies [30,33,35]. Therefore, a dosing interval similar to TAK-242 was adopted to ensure a consistency in injections and animal handling across groups.

### Ex vivo electrophysiology

One-week after FPI or sham injury, animals were anesthetized with isoflurane and decapitated. Brain was quickly harvested, hemisected and coronal hippocampal slices (300 µm) were prepared in ice-cold sucrose artificial cerebrospinal fluid (aCSF) containing (in mM): 85 NaCl, 75 sucrose, 24 NaHCO3, 25 glucose, 4 MgCl2, 2.5 KCl, 1.25 NaH2PO4, and 0.5 CaCl2 [36]. Slices were incubated at 32°C for 20 min in an interface holding chamber containing an equal volume of sucrose-aCSF and recording aCSF containing (in mM) 126 NaCl, 2.5 KCl, 2 CaCl_2_, 2 MgCl_2_, 1.25 NaH_2_PO_4_, 26 NaHCO_3_ and 10 D-glucose. All solutions were saturated with 95% O_2_ and 5% CO_2_ and maintained at a pH of 7.4. Slices were allowed to equilibrate for an hour at room temperature before recordings. Whole-cell voltage- and current-clamp recordings were obtained from dentate granule cells (DGCs) in the suprapyramidal blade at 33-34°C under IR-DIC (Olympus BX51 with 40x water immersion objectives) using 3-6MΩ glass pipettes fabricated using dual stage glass micropipette puller (Narishige PC-10, Amityville NY). Data was recorded using MultiClamp 700B amplifiers, digitized using Digidata 1440A and acquired using pClamp10 (Molecular Devices, San Jose, CA) at 10-kHz sampling frequency. Spontaneous and miniature excitatory/inhibitory currents from the same DGCs were obtained using a cesium-based internal solution (in mM: Cesium methanesulfonate 140, NaCl 5, HEPES 10, EGTA 0.2, Mg2ATP 2, Na2GTP 0.2, Qx314 5, (pH 7.25, 270-285 mOsm). Biocytin (0.2%) was included in the internal solution for post hoc cell identification. Spontaneous and miniature excitatory postsynaptic potentials (sEPSC and mEPSC) were recorded from the same neurons at a holding potential of −70 mV, and spontaneous and miniature inhibitory postsynaptic potentials (sIPSC and mIPSC respectively) were recorded at a holding potential of 0 mV. Action potential independent miniature currents were isolated using tetrodotoxin (TTX, 1μM). In a subset of experiments TLR4 signaling was selectively activated by incubating slices in LPS-B5 ultrapure (Escherichia Coli serotype O55:B5, 10 µg/ml, Invivogen, San Diego, CA), a selective TLR4 agonist, for 60-90 minutes before recording [37]. Data from cell in which access resistance changes over 20% during recordings and recordings with access resistance above 25 MΩ were excluded from analysis. Spontaneous/miniature synaptic currents (100 events/cell) were detected using template search feature in Easy Electrophysiology (version 2.6., London, UK.) by a blinded investigator and interevent intervals and amplitudes were analyzed as cumulative distribution in Graphpad Prism 10.0.1 (Boston, MA).

### In vivo electrophysiology

One-week after FPI/Sham injury rats were anesthetized under urethane (1.5g/kg i.p.) and head-fixed onto a stereotaxic frame. Under aseptic conditions, rats were implanted with a 50 µm tungsten wire depth electrode (California Fine Wire company, Grover beach, CA) in the granule cell layer (AP: 3-4 mm, ML: 2 mm, DV: 3-3.5 mm) guided by electrical stimulation of the angular bundle (ML:4-4.2 mm, DV: 1.8-2 mm; pulse width 150µs delivered at 0.1Hz). Two screw electrodes on the contralateral side served as reference and ground. The position of electrodes was secured using dental cement. Acute recordings from urethane anesthetized rats started 60 minutes after induction. Current-response curves were obtained in response to electrical stimulation of the angular bundle (150 µs @ 0.1Hz). For assessing Long Term Potentiation (LTP), DG LFP responses were evoked every 30 seconds for a period of 30 minutes to establish a baseline. LTP was induced by Theta Burst Protocol consisting six trains of six pulses at 400 Hz, 200 ms between trains, delivered six times at 20 seconds interval [38] and responses were recorded for an additional 180 minutes. At the end of the experiment, animals were euthanized with a lethal dose of Euthasol (150 mg/kg, i.p.).

Signals were amplified 100 times (Pinnacle systems, Lawrence, KS), sampled at 10kHz, digitized using PowerLab 16/35 and recorded using LabChart8 software (AD Instruments, Colorado Springs, CO). Analysis was conducted using LabChart modules. Dentate spikes were recorded from urethane anesthetized rats (1.5g/kg i.p.) using 16 Ch silicon probe (A1x16-5mm-100-703 CM16, Neuronexus, Ann Arbor, MI). Signals were amplified 100 times using Model 3600 amplifier (AM-Systems, Sequim, WA), digitized using PowerLab 16/35 and recorded using Labchart8 Software (AD instruments, Colorado Springs, CO). Dentate spikes were detected using Toothy analysis pipeline[39] (https://github.com/Farrell-Laboratory/Toothy). Briefly, large positive events were identified using a 4.5 S.D. threshold. Detected events were visually confirmed by a blinded investigator and the total number of dentate spikes recorded in a 30-minute period were counted. We did not classify dentate spikes as type 1 or type 2.

### In Situ Zymography

MMP-9 activity was evaluated in hippocampal slices obtained from rats and mice 48 hours after injury using a modified in situ zymography protocol [40]. Briefly, sham/injured rats or TLR4^f/f^:CaMKIIa-CreERT2 mice and Cre^-ve^ controls were transcardially perfused using an alcohol-based formalin free tissue fixative (Accustain, A5472 Sigma-Aldrich, St.Louis, MO), embedded in 2% agar and coronal slices (50μm, VF-310-0Z Compresstome, Precisionary Instruments, Ashland, MA) were prepared at <4°C to preserve enzyme activity. Slices were then incubated with a fluorogenic Dye Quenched gelatin substrate (1:50 dilution, Invitrogen, Waltham MA) for 40 min at 37°C, washed with PBS, fixed in 4% paraformaldehyde and mounted on slides in Vectashield (Vector Labs, Newark, CA). The specificity of reactions was confirmed by adding MMP-9 inhibitor SB-3CT to control slices (see supplementary figure 2J-K). Semiquantitative analysis of DQ fluorescence in DG molecular layer were performed using ImageJ by a blinded investigator. Data was collected from 2-4 slices / animals and averaged. Intensity measures were normalized to CA1 stratum radiatum since we did not observe any effect of injury, treatment or genotype in this region (Supplementary figure 4I-J).

### Single molecule fluorescent in situ hybridization (RNAscope)

To visualize TLR4 in mice and MMP-9 mRNA in rat, we used RNAscope Multiplex Fluorescent in situ hybridization on cryosectioned slices (20μm). RNAscope Multiplex Fluorescent Reagent kit v2 assay (ACD, Newark, CA) was used following the protocol for fixed tissue [41]. The following kits were used for this study: RNAscope Multiplex Fluorescent Reagent kit (Cat#323100), RNAscope Probe-MM-TLR4 (Cat#316801) and RNAscope probe-Rn-MMP-9 (Cat#532281). Slices were coverslipped with DABCO with DAPI mounting media and imaged using a Zeiss Axioscope and analyzed using QuPath software by a blinded investigator.

### Immunohistochemistry

Male and female TLR4^f/f^:CaMKIIa-CreERT2 mice (8-10 weeks) were induced with tamoxifen as described above and subjected to Sham or FPI. Three days after injury, brain tissue was harvested following transcardial perfusion with PBS followed by 4% PFA. Brains were post-fixed overnight and then transferred to 30% sucrose until fully equilibrated. Tissue was cryosectioned at 20 µm thickness and mounted on Superfrost Plus glass slides. For immunohistochemistry, slides were brought to room temperature to remove OCT and washed three times in PBS. Sections were permeabilized with 0.2% Triton X-100 in PBS for 45 minutes at room temperature, followed by blocking with 20% goat serum for 45 minutes. Sections were then incubated overnight at 4 °C with a primary antibody against TLR4 (mouse anti-TLR4; Santa Cruz Biotechnology; 1:250). After three PBS washes, sections were incubated with goat anti-mouse Alexa Fluor 546 secondary antibody (1:1000, Thermo Fisher, S11227) for 1 hour at room temperature. Slides were washed three times with PBS and mounted using an antifade mounting medium. Images were acquired using a Zeiss Axioscope-5 with stereo investigator (MBF Bioscience) from three animals per group, with six sections analyzed per animal. Quantification was performed using ImageJ software by calculating the Corrected Total Cell Fluorescence (CTCF) in the granule cell layer and hilus for each section (Supplementary figure 3 A-C).

### Stereological Estimation of Hilar Neurons

To quantify neuronal loss following injury, sham and FPI rats (n = 3 and 4 rats, respectively., 1-month post-injury) were deeply anesthetized with Euthasol (150 mg/kg, i.p.) and transcardially perfused with PBS followed by 4% PFA. Brains were post-fixed and coronally sectioned (50 µm) using a compressotome (VF-310-Z, Precisionary Instruments). Sections from the side of injury were processed for Nissl staining (0.1% cresyl violet acetate and 0.25% glacial acetic acid). Unbiased stereological estimates of hilar neuron populations were obtained using the Optical Fractionator probe (Stereo Investigator; MBF Bioscience) on a Zeiss Axioscope-5 microscope by blinded investigators (See Supplementary Figure 1). Cell counts from three slices were averaged within each animal and used for statistical analysis.

### Behavioral Experiments

Rats were trained on Barnes Maze task to assess spatial learning 1-month post-injury. The Barnes maze table (Maze Engineers, https://conductscience.com/maze/) was 122 cm in diameter with 20 holes (10 cm each). The target hole was equipped with an escape box (22.5 cm x 14 cm x 9 cm LxWxH). The maze table was set up within black curtain walls with visual cues and a camera installed above to record animal movement. Two bright floodlights and a fan served as aversive stimuli to encourage animals reach the escape box. Animals were held in their home cage outside the curtain in a dark room until their turn to run the trial. On day 1, rats were habituated to the behavior room for at least 1 hour and then placed in the middle of the table inside starter cup for 1 minute following which they were allowed to explore the table for 180 seconds. At the end of the trial, animals were gently guided to their designated escape hole. The escape hole locations were randomly assigned to each rat and remained the same till the end of the experiment. The table was thoroughly wiped with ethanol between tests to eliminate odor cues. Rats were subjected to a single 180-second trial for 3 days. Task acquisition and retention were confirmed through tests on day 4 and day 7. Four rats that fell from the table and exhibited signs of lethargy were removed from the test and sacrificed. Behavioral analysis was performed by a blinded experimenter using Any-maze software. Parameters such as latency to escape hole, distance travelled and spatial search strategy were evaluated between sham and injured rats. Spatial search strategies were scored based on 11-point scoring scheme[42]. Thigmotaxis was quantified by defining a peripheral zone of the testing arena that was approximately 12 cm from the outer edges of the apparatus. The proportion of time the animals spent within this designated zone was recorded and used as the metric for anxiety-like behavior [43].

### Statistics

Data are presented as mean ± standard error of the mean (SEM) for parametric measures, or as median and interquartile range (IQR) where non-parametric distributions were observed. To ensure rigorous data interpretation and minimize investigator bias, all data analysis, including quantification of in situ zymography, electrophysiological recordings, and behavioral scoring, was performed by researchers blinded to the treatment groups and genotypes.

Normality of data distributions was assessed using the Shapiro-Wilk test. For the analysis of synaptic events (sEPSC, mEPSC, sIPSC, and mIPSC), cumulative probability distributions of inter-event intervals (IEI) and amplitudes were compared across groups using equal number of consecutive events with the non-parametric Kolmogorov-Smirnov (K-S) test. Effect sizes for these distributions were calculated using Cohen’s d. When multiple comparisons were involved (Figures 1&3, Supplementary figure 2) Holms-Bonferroni correction was adopted and corrected *P* values are reported (See Supplementary table 3). Differences between two groups were evaluated using two-tailed unpaired t-tests. For experiments involving multiple factors, such as injury and drug treatment or injury and genotype, two-way ANOVA, two-way repeated measures (RM) ANOVA (for input/output curves and behavioral training days) or three-way ANOVA were employed, followed by Šídák’s multiple comparisons test where appropriate. Significance was defined at P<0.05. Statistical outlier tests were not applied. Statistical analyses were performed using Graphpad Prism 10.0.1 (Boston, MA).

**Figure 1:**
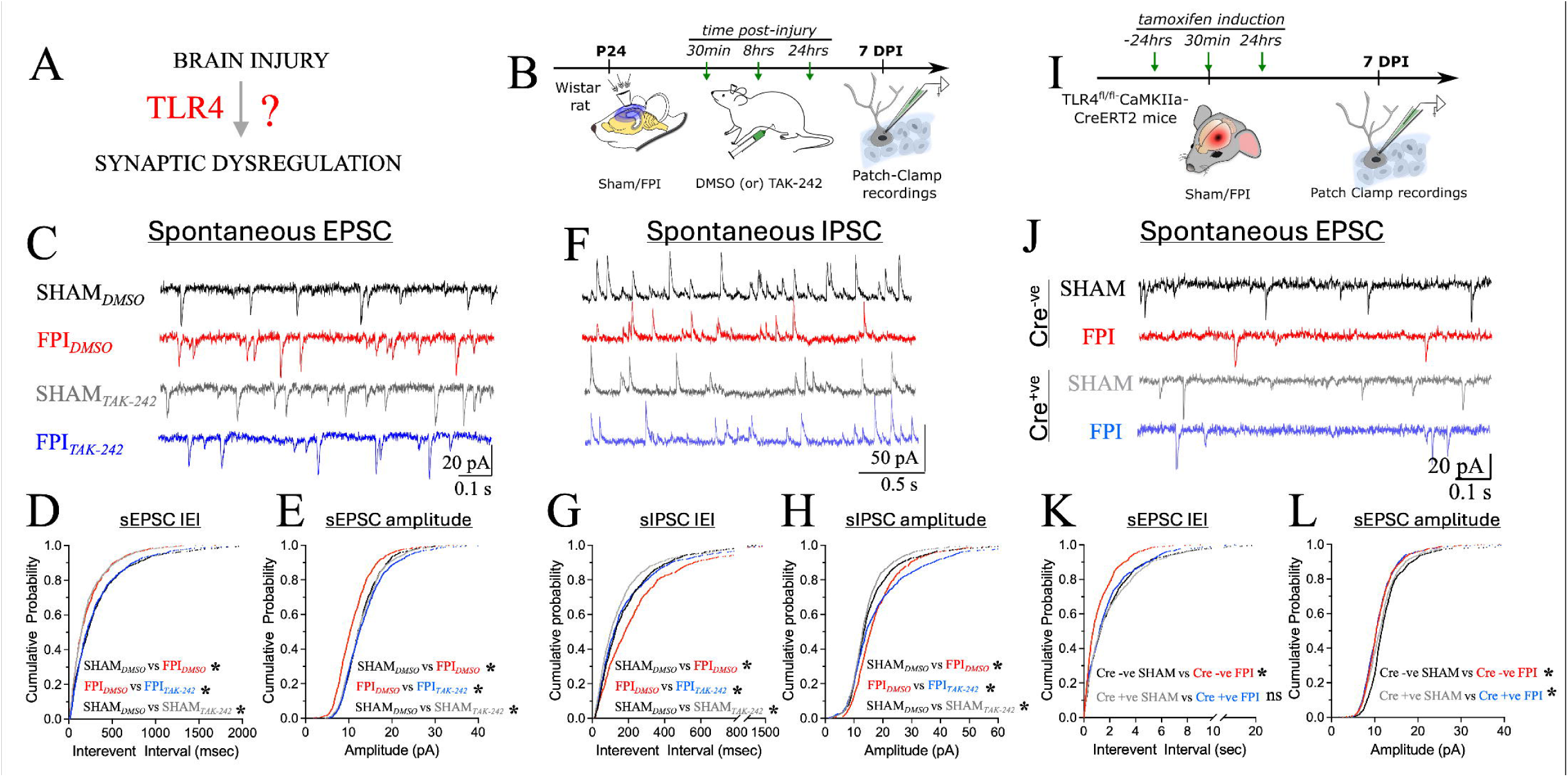
TLR4 signaling drives early excitatory and inhibitory synaptic changes in DGCs after brain injury. A) Schematic of the experimental question. B) Experimental timeline for examining synaptic changes after FPI in rats. C) Representative sEPSC traces showing whole-cell voltage clamp recordings from granule cells in vehicle treated sham controls (black), vehicle treated FPI (red), TLR4 inhibited sham (grey) and TLR4 inhibited FPI (blue). D-E) Cumulative probability plots of sEPSC interevent intervals (D) and amplitude (E) in vehicle treated and TLR4 inhibited sham and injured rats. F) Representative sIPSC traces showing whole-cell voltage clamp recordings from granule cells in vehicle treated sham controls (black), vehicle treated FPI (red), TLR4 inhibited sham (grey) and TLR4 inhibited FPI (blue). G-H) Cumulative probability plots of sIPSC interevent intervals (G) and amplitude (H) in vehicle treated and TLR4 inhibited sham and injured rats. I) Experimental timeline for examining synaptic changes after FPI in TLR4^fl/fl^: CaMKIIa-CreERT2 mice. J) Representative sEPSC traces from granule cells in Cre^-ve^ sham controls (black), Cre^-ve^ FPI (red), Cre^+ve^ sham (grey) and Cre^+ve^ FPI (blue). K-L) Cumulative probability plots of sEPSC interevent intervals (K) and amplitude (L) in granule cells from Cre^-ve^ and Cre^+ve^ mice. n = 8-9 cells from 3 rats/group (A-H) and 7 cells from 3 mice/group (I-L). * indicates *P*<0.05 by K-S test with Holm-Šídák correction.

## RESULTS

### Neuronal TLR4 signaling is necessary for early excitatory circuit changes after brain injury

Prior studies have identified TLR4 contribution to early changes in glutamatergic synapse strength after brain injury [12–14]. However, the nature of early circuit reorganization after brain injury in the dorsal hippocampus and whether immune signaling regulates these changes has not been examined. Whole-cell voltage clamp recordings from DGCs in dorsal hippocampal slices one-week after FPI revealed an early decrease in the inter-event intervals (IEI) of spontaneous excitatory postsynaptic currents (sEPSCs), indicating an increase in sEPSC frequency (*P*<0.0001 by K-S test with Holm-Šídák correction; Cohen’s *d* = 0.34, *n* = 8 cells/ group, Figure 1A-E, Supplementary table1 & 3). Additionally, we observed a reduction in sEPSC amplitude in DGCs from rats after brain injury (*n* = 8 cells/ group, *P*<0.0001 by K-S test, Cohen’s *d* = 0.41). To assess potential post-traumatic synaptic reorganization, we recorded action potential-independent miniature EPSCs (mEPSCs) in DGCs the presence of tetrodotoxin (TTX, 1 μM). Similar to sEPSCs, DGCs in rats one week after FPI exhibited shorter mEPSC IEI and smaller mEPSC amplitudes compared to Sham controls (*P* <0.001 by K-S test with Holm-Šídák correction; Cohen’s *d* = 0.22 for IEI and 0.85 for amplitude, *n* = 7 cells/ group, Supplementary figure 2A-C, Supplementary table 2 & 3). Together, these findings provide the first evidence of an increase in excitatory synaptic inputs to DGCs in dorsal hippocampus of rats one week after injury.

Significant changes in synaptic inhibition are often observed early after brain injury [5,7,44]. To evaluate whether inhibitory circuits are reorganized alongside the excitatory circuits, we examined spontaneous inhibitory postsynaptic potentials (sIPSC) in the same neurons. In contrast to changes in sEPSCs, sIPSC IEI was prolonged in DGCs from injured rats (*P* <0.0001 by K-S test with Holm-Šídák correction; Cohen’s *d* = 0.29, n= 8 cells/group, Figure 1F-H, Supplementary tables 1 & 3). These findings indicate that in addition to receiving more frequent excitatory inputs, the DGCs receive fewer inhibitory inputs after brain injury. DGCs also showed an increase in sIPSC amplitude compared to sham controls after brain injury (*P* <0.0001 by K-S test with Holm-Šídák correction; Cohen’s *d* = 0.16, *n* = 8 cells/ group). These findings differ from our findings in the ventral hippocampus demonstrating an increase in sIPSC frequency in DGCs after FPI [36,45,46]. We also observed an increase in mIPSC amplitudes with no change in mIPSC IEI (Supplementary Figure 2D-F, Supplementary tables 2 & 3). Together, these findings indicate an early dysregulation in excitatory and inhibitory inputs to DGC’s one-week after FPI. Furthermore, these results identify early circuit reorganization after FPI and reveal potential differences in synaptic inhibition and post-traumatic plasticity in the dorsal vs ventral hippocampus.

To determine whether inhibition of TLR4 signaling early after injury modified DGC synaptic currents, we treated sham and FPI rats with TLR4 inhibitor TAK-242 (0.5mg/kg, i.p, 30 min, 8 hours and 24 hours post-injury). TAK-242 treatment reduced changes in sEPSC and mEPSC frequency and amplitude in injured rats (*P* <0.0001 by K-S test with Holm-Šídák correction; Cohen’s *d* = 0.62 for sEPSC IEI, 0.50 for sEPSC amplitude, 0.54 for mEPSC IEI and 0.89 for mEPSC amplitude, *n* = 8-9 cells/ group, Figure.1A-E, Supplementary figure 2A-C, and Supplementary tables 1 & 3). In addition, TAK-242 reduced injury-driven changes in sIPSC frequency and amplitude (*P* <0.0001 by K-S test with Holm-Šídák correction; Cohen’s *d* = 0.22 for IEI, 0.11 for amplitude, *n* = 7-8 cells/ group, Figure.1F-H, Supplementary tables 2 & 3). While mIPSC frequency was unaltered, mIPSC amplitude was reduced by TAK-242 treatment in injured rats (*P* <0.0001 by K-S test with Holm-Šídák correction; Cohen’s *d* = 0.47 for amplitude, *n* = 6-7 cells/ group, Supplementary figure 2D-F, Supplementary tables 2 & 3). These findings demonstrate a role for TLR4 signaling in driving early synaptic dysregulation after FPI.

In contrast to injured animals, sham controls receiving TAK-242 showed paradoxically opposing changes in excitatory and inhibitory synaptic inputs to DGCs. In sham controls, TAK-242 treatment reduced sEPSC IEI in DGCs, increasing frequency as observed after FPI but failed to alter sEPSC amplitude (*P* <0.0001 by K-S test with Holm-Šídák correction; Cohen’s *d* = 0.35 for IEI, *n* = 8-9 cells/ group, Figure 1B-E). Both mEPSC interval and amplitude were reduced in TAK-242 treated sham controls (*P* <0.0001 by K-S test with Holm-Šídák correction; Cohen’s *d* = 0.24 for IEI and 0.30 for amplitude, *n* = 7-8 cells/ group, Supplementary figure 2A-C, Supplementary tables 2 & 3) consistent with synaptic reorganization. Moreover, TAK-242 treated sham controls showed a reduction in DGC sIPSC IEI and amplitude suggesting more frequent inhibitory inputs (*P* <0.01 by K-S test with Holm-Šídák correction; Cohen’s *d* = 0.07 for IEI and 0.21 for amplitude, *n* = 7-8 cells/ group, Figure 1F-H, Supplementary tables 1 & 3). However, TAK-242 treatment did not alter mIPSC frequency or amplitude. Together, these findings identify a novel contribution of basal TLR4 signaling in maintaining excitatory synaptic homeostasis in uninjured animals.

Our previous studies demonstrated neuronal TLR4 signaling as critical for post-traumatic increase in DGCs AMPA currents [13]. To test whether the glutamatergic circuit changes identified above require neuronal TLR4 signaling, we examined changes in sEPSC frequency and amplitude in mice with selective deletion of TLR4 in CaMKII expressing excitatory neurons (TLR4^fl/fl^-CaMKIIa-CreERT2 mice). Care was taken to avoid constitutive and developmental effects by inducing Cre recombinase just prior to injury.

Immunostaining for TLR4 in mice three-days after FPI, following the 3-day tamoxifen induction strategy, revealed strong immunostaining for TLR4 in the granule cell layer and hilus of Cre^-ve^ mice which was absent in Cre^+ve^ mice, confirming TLR4 deletion in DGCs (Nested t-test: t(4) = 7.57, *P* = 0.0016, F(1,4) = 58.82, *n* = 5-6 slices from 3 mice/ group, Supplementary figure 3A-C). In addition, RNAscope analysis of TLR4 mRNA confirmed the lack of injury-driven TLR4 mRNA increase in Cre^+ve^ mice (F(2,9) = 8.353, *P*<0.05 by one-way ANOVA, *n* = 3-5 mice/ group, Supplementary figure 3D-F). Validation studies to test for effect on TLR4 deletion on neuronal intrinsic physiology failed to reveal genotype-dependent effects on the resting membrane potential, input resistance or rheobase in DGCs from sham controls (Supplementary figure 4C-E). However, there was a significant decrease in capacitance and an increase in firing frequency in Cre^+ve^ sham mice compared to Cre^-ve^ sham mice (Capacitance - Effect of Genotype-F(1,23) = 6.9, *P* = 0.015 by two-way ANOVA; Firing frequency – Effect of Genotype – F(1,23) = 4.706, *P* =0.040 by three-way ANOVA, *n* = 7 cells/ group, Supplementary figure 4B&F). Consistent with our previous reports in rat and mice [31,47], there were no injury effects on DGC active properties in either genotype. While there was no injury effect on DGC resting membrane potential in Cre^-ve^ mice, DGCs from Cre^+ve^ mice with TLR4 deletion were more depolarized (Supplementary figure 4C). Together these validation studies demonstrate the lack of injury-induced TLR4 increase in Cre^+ve^ mice and suggest a role for TLR4 in basal DGC excitability and maintenance of RMP after injury.

Having established the lack of posttraumatic TLR4 upregulation in Cre^+ve^ mice we examined the contribution of neuronal TLR4 signaling to injury-induced changes in synaptic inputs to DGCs. Similar to our findings in rat, brain injury reduced DGC sEPSC IEI and amplitude in Cre^-ve^ mice with intact TLR4 signaling (*P* <0.0001 by K-S test with Holm-Šídák correction; Cohen’s *d* = 0.42 for IEI and 0.31 for amplitude, *n* = 7 cells/ group, Figure 1I-L, Supplementary table 1). Importantly, DGCs from Cre^+ve^ mice with conditional deletion of TLR4 in excitatory neurons failed to exhibit injury-induced decrease in sEPSC IEI, but showed a decrease in sEPSC amplitude after brain injury (*P* = 0.0008 by K-S test with Holm-Šídák correction; Cohen’s *d* = 0.16 for amplitude, *n* = 7 cells/ group. Interestingly, DGC sEPSC IEI and amplitude were not different between Cre^-ve^ and Cre^+ve^ sham injured mice. These findings demonstrate that TLR4 expression in glutamatergic neurons is necessary for the early injury-driven changes in excitatory synaptic inputs to DGCs.

### MMP-9 activity is required for TLR4-dependent changes in DGC excitatory synapses after brain injury

We next sought to identify downstream molecular mechanisms through which TLR4 signaling dysregulates DGC synaptic inputs after brain injury. Several studies have reported an upregulation in Matrix Metallproteinase-9 (MMP-9) expression in pre-clinical models of TBI [27,30,35]. MMP-9 is a critical regulator of synaptic strength and plasticity [17,18], and perturbations in MMP-9 activity is known to dysregulate synaptic physiology [19,40,48–51]. Although extrinsic activation of TLR4 signaling has been shown to enhance MMP-9 expression in primary astrocyte/neuronal co-cultures and *in vivo* [15,16], whether TLR4 regulates MMP-9 activity and if this can induced synaptic and functional changes *in vivo* and in the injured brain remains to be determined. Moreover, whether *neuronal* TLR4 regulates MMP-9 activity has not been examined.

To directly assess MMP-9 activity, we performed *in-situ* zymography on hippocampal slices obtained from sham/FPI rats 48 hours after injury, focusing on the DG molecular layer where the afferent glutamatergic synapses are located. DQ-fluorescence intensity values were normalized to CA1 Stratum Radiatum (SR) since no changes were observed in this region with injury, treatment or genotypes (Supplementary figure 4I-J, see methods). Our experiments identified a significant increase in the MMP-9 activity in the DG molecular layer in brain-injured rats compared to sham controls (MMP-activity normalized to CA1 SR: Sham:14.56±2.15%, FPI:30.43±2.89%, *n* = 5 sham and 6 FPI rats, Figure 2A-C).

**Figure 2:**
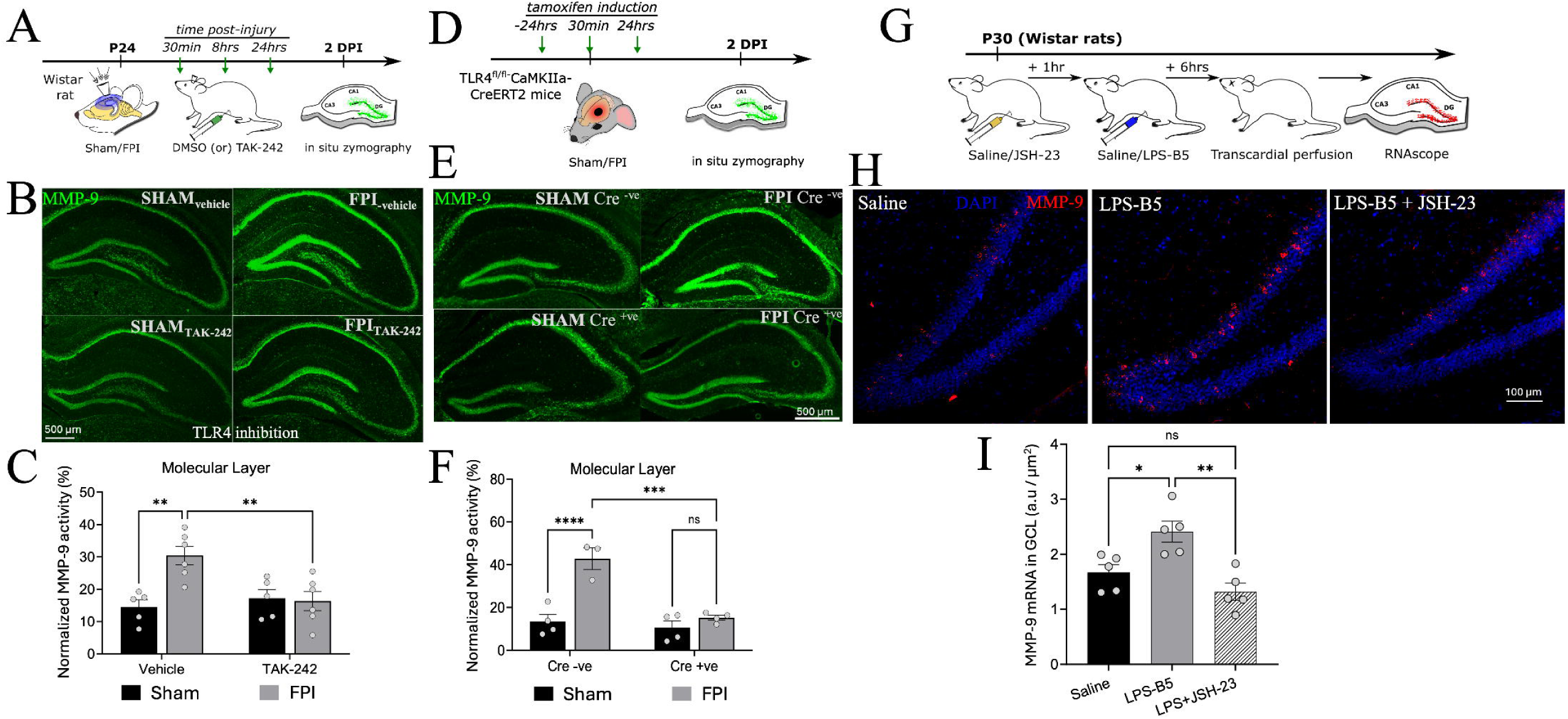
TLR4 signaling upregulates MMP-3 activity in the dentate gyrus after brain injury. A) Experimental timeline for in-situ zymography to quantify MMP-9 activity after brain injury in rat. B) Representative images showing MMP-9 activity in hippocampal sections from sham and injured rats. Images from vehicle treated (middle) and TLR inhibited mice (bottom) show MMP-9 activity in green. C) Quantification of MMP-9 activity, normalized to CA1 stratum Radiatum, shows an increase in MMP-9 activity in vehicle treated injured rats, but not in TAK-242 treated injured rats. Note that MMP-9 activity in vehicle treated and TAK-242 treated sham rats were not different. D) Experimental timeline for in-situ zymography to quantify MMP-9 activity after brain injury in mice. E) Representative images showing MMP-9 activity in sham and injured mice. Images from hippocampal sections from Cre^-ve^ sham and injured mice (top) and Cre^+ve^ sham and injured mice (bottom) show MMP-9 activity in green. F) Quantification of MMP-9 activity, normalized to CA1 stratum Radiatum, shows an increase in MMP-9 activity after brain injury in Cre^-ve^ mice, but not in Cre^+ve^ mice. Note that MMP-9 activity in sham mice was not different between genotypes. * indicates *P*<0.05, ** indicates *P*<0.01, *** indicates *P*<0.001, **** indicates *P*<0.0001 by TW-ANOVA with Šídáks multiple comparisons post hoc tests. G) Experimental timeline for RNAscope experiments in naïve rats. H) Representative images showing increase in MMP-9 mRNA in sections from rats after LPS-B5 treatment. I) Summary plot of mRNA levels in saline, LPS-B5 treated and LPS-B5+JSH-23 sections shows an increase in mRNA levels after LPS-B5 treatment which is abolished by blocking NF-κB. GCL-granule cell layer. n = 5-6 rats/group (A-C), 3-4 mice /genotype (D-F), and 5 rats/group (G-I). * indicates *P*<0.05, ** indicates *P*<0.01, by OW-ANOVA with Šídáks multiple comparisons post hoc tests. Data points in panel C, F and I represent individual animals.

Pharmacologically blocking TLR4 early after FPI reduced MMP-9 activity in the DG of brain injured rats (Sham_TAK-242_: 17.22±2.68%, FPI _TAK-242_: 16.36±2.97%, *n* = 5 Sham and 6 FPI rats; Effect of Injury x Treatment F (1,18) = 9.13; *P* <0.05 by Two-way ANOVA). In contrast, MMP-9 activity in sham controls treated with TAK-242 was not different from vehicle treated sham controls. These results identify TLR4 regulation of MMP-9 activity in the injured brain.

To isolate the role of neuronal TLR4 in regulating MMP-9 activity, we performed in-situ zymography on mice with neuronal TLR4 deletion (TLR4^fl/fl^-CaMKIIa-CreERT2) and Cre^-ve^ controls (Supplementary figure 4) 48 hours after Sham or FPI. We did not observe any difference in MMP-9 activity between Cre^-ve^ sham and Cre^+ve^ sham mice, indicating that TLR4 deletion does not alter normal MMP-9 levels without injury (Figure 2D-F). While Cre^-ve^ mice showed an increase in MMP-9 levels after injury, mice with selective TLR4 deletion in glutamatergic neurons failed to develop elevated MMP-9 activity after FPI (MMP-activity normalized to CA1 SR: Cre^-ve^ Sham: 13.43±3.36%, Cre^-ve^ FPI: 42.79±5.02%; Cre^+ve^ SHAM: 10.59±3.15%, Cre^+ve^ FPI: 12.20±1.16%., *n*=3-4 mice/group; Effect of Injury x Genotype, F (1,11) = 14.70; *P*<0.05 by Two-way ANOVA, Figure 2D-F). These findings demonstrate that neuronal TLR4 is necessary for the post-injury upregulation of MMP-9 activity.

Next, we tested the possibility that TLR4 signaling increased MMP-9 levels through transcriptional upregulation. Naïve rats (P30) were injected with a selective TLR4 agonist, LPS-B5 (5mg/kg., i.p.) or saline and euthanized after six hours for single molecule fluorescent in situ hybridization (smFISH) to quantify MMP-9 mRNA levels. Sections from LPS-B5 injected rats showed a significant increase in MMP-9 mRNA in DGCs, indicating that TLR4 activation enhances MMP-9 via transcriptional upregulation (intensity A.U/μm^2^: saline controls 1.67±0.15; LPS-B5 treated 2.41±0.19; n= 5/group, Figure 2G-I). Furthermore, inhibition of NF-κB, a key downstream effector of TLR4, using JSH-23 (10 mg/kg) administered one hour before LPS-B5 injection abolished the upregulation of MMP-9 (intensity A.U/μm^2^: JSH-23 + LPS-B5 1.319±0.16, n = 5 rats, Main effect F(2,12) =11.20, *P* = 0.002, Figure 2G-I) Together, these findings identify TLR4 and NF-κB dependent transcriptional upregulation of MMP-9 as a candidate mechanism for increase in MMP9 activity in the DG following brain injury.

MMP-9 has been shown to play a crucial role in plasticity of excitatory and inhibitory synapses [52]. Therefore, we reasoned that TLR4 mediated increase in MMP-9 activity may be required for the post-traumatic changes in synaptic inputs to DGCs. To examine the acute effects of TLR4 signaling *ex vivo*, we compared sEPSCs in hippocampal slices from sham rats incubated in LPS-B5 (10 μg/ml for 90min) or aCSF. Acute LPS-B5 incubation significantly reduced sEPSC IEI in DGCs, mimicking the increase in sEPSC frequency observed in FPI animals (Sham vs LPS-B5 incubated: *P* <0.0001 by K-S test with Holm-Šídák correction; Cohen’s *d* = 0.20 for IEI, *n* = 8 cells from 4 rats, Figure 3A-B,C). However, LPS-B5 incubation increased the sEPSC amplitude (Figure 3A-B,D). Importantly, inhibiting MMP-9 using SB-3CT (10 μM) mitigated LPS-B5-induced changes in sEPSC IEI indicating that TLR4-mediated alterations in excitatory synaptic inputs to DGCs require MMP-9 activity (LPS-B5 incubated vs SB-3CT+LPS-B5: *P* <0.0001 by K-S test with Holm-Šídák correction; Cohen’s *d* = 0.21 for IEI, *n* = 9 cells from 4 rats, Figure 3A-C). Conversely, SB-3CT co-incubation failed to reduce, and further exacerbated, the LPS-B5 mediated increase in sEPSC amplitude (Figure 3B, D). Acute TLR4 activation failed to alter sIPSC frequency but increased mIPSC interevent intervals in DGCs (Supplementary figure 5A-B).

**Figure 3:**
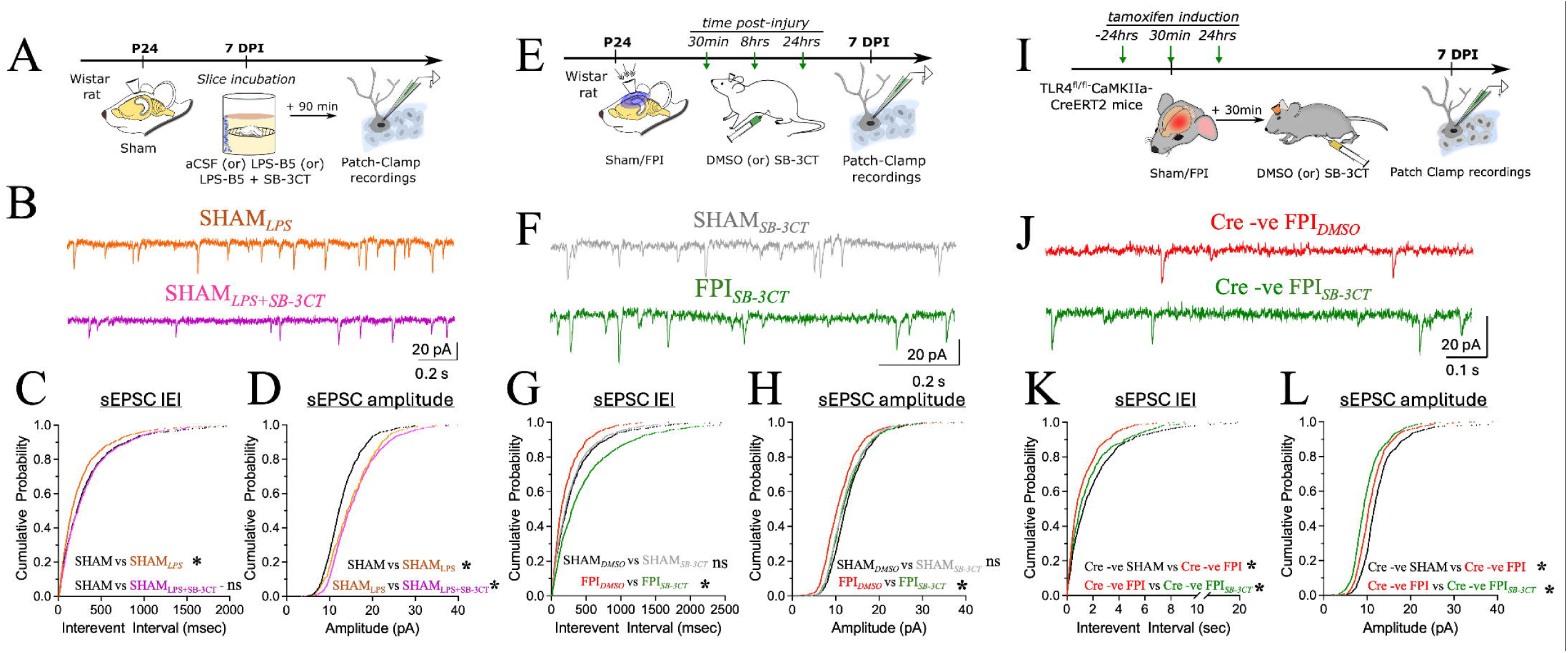
Blocking MMP-3 prevents early synaptic changes in excitatory inputs to DGCs after brain injury. A) Experimental timeline for examining synaptic changes in acute rat slices after LPS incubation. B) Representative sEPSC traces from granule cells in LPS incubated (orange), LPS+SB-3CT co-incubated (magenta) slices. C-D) Cumulative probability plots of sEPSC interevent intervals (C) and amplitude (D) between LPS-B5 incubated and LPS-B5+SB-3CT co-incubated slices. Note that LPS incubation significantly reduced interevent intervals. Co-incubation of slices in MMP-9 blocker SB-3CT prevented this change. E) Experimental timeline for examining synaptic changes in SB-3CT treated sham/injured rats. F) Representative granule cell sEPSC traces showing from SB-3CT treated sham (grey) and injured rats (green). G-H) Cumulative probability plots of sEPSC interevent intervals (G) and amplitude (H) in SB-3CT treated sham/injured rats. I) Experimental timeline for examining synaptic changes in SB-3CT treated Cre^-ve^ mice after brain injury. J) Representative granule cell sEPSC traces from vehicle treated (red) and SB-3CT treated mice(green). K-L) Cumulative probability plots of sEPSC interevent intervals (K) and amplitude (L) in vehicle treated and SB-3CT treated Cre^-ve^ mice. n = 8-9 cells(A-D), 7-10 cells (E-H) from 3-4 rats/group and 7 cells from 3 mice /group (I-L). * indicates *P*<0.05 by K-S test with Holm-Šídák correction.

Since we find that TLR4-mediated alterations in DGC sEPSC frequency requires MMP-9 activity, we tested if systemic inhibition of MMP-9 using SB-3CT treatment (50 mg/kg, i.p.) early after FPI would prevent post-traumatic changes in excitatory inputs to DGCs. SB-3CT treatment effectively mitigated injury-induced the changes in sEPSC IEI and amplitude one-week after FPI in rat (*P* <0.0001 by K-S test with Holm-Šídák correction; Cohen’s *d* = 0.60 for IEI and 0.30 for amplitude, *n* = 7 cells / group, see Supplementary table1&3, Figure 3E-H). Since Cre^+ve^ mice failed to show posttraumatic increase in sEPSC frequency (Figure 1K), we examined Cre^-ve^ mice to assess the role of MMP9 in injury-induced changes in sEPSC in mice. Consistent with our findings in rat, SB-3CT treatment reduced post-injury changes in sEPSC IEI in DGCs from Cre^-ve^-negative TLR4^fl/fl^-CaMKIIa-CreERT2 mice (*P* <0.0001 by K-S test with Holm-Šídák correction; Cohen’s *d* = 0.283 for IEI, *n* = 9 cells Figure 3I-L). SB-3CT treatment did not have any effect on sIPSC frequency or amplitude (Supplementary figure 5G-I) after FPI suggesting that TLR4 regulation of DGC inhibitory synaptic parameters after injury does not require MMP-9 activation. However, SB-3CT caused a small increase in sIPSC amplitude in sham controls (Supplementary figure 5I). Together, these results indicate that injury-driven changes in excitatory, but not inhibitory, inputs to DGCs required both TLR4 signaling and MMP-9 activity.

### Early activation of TLR4 - MMP-9 molecular axis contributes to DG network dysfunction after brain injury

Having established that the TLR4-MMP-9 mechanistic axis increases glutamatergic inputs to DGCs after brain injury, we examined the contribution of this molecular pathway to early changes in DG circuit function in rats *in vivo* one-week after brain injury. Input/output plots of the DG population spike amplitude in response to perforant path stimulation at increasing current intensities was recorded under urethane anesthesia. DG population spike responses normalized to peak amplitude, revealed a leftward shift after FPI indicating increased network excitability (*n* = 6 SHAM and 5 FPI rats; Effect of injury – F (1,9) = 6.462, *P =* 0.031 by Two-way RM ANOVA, Figure 4A-C). This increase in network excitability was mitigated by early inhibition of either TLR4 signaling or MMP-9 activity, demonstrating their contribution in promoting post-traumatic DG circuit dysfunction (n= 5-6 rats/group; Effect of Stimulus x Treatment: F (6,166,40.08) = 4.752, *P*=0.0009 by Two-way RM ANOVA, Figure 4D). Paradoxically, as previously reported in slices [12–14], TAK-242 treatment increased DG population spike amplitude in sham controls (Figure 4E). However, consistent with its lack of effect on EPSC frequency in sham controls, SB-3CT treatment did not alter DG network excitability in sham rats (Figure 4E).

**Figure 4:**
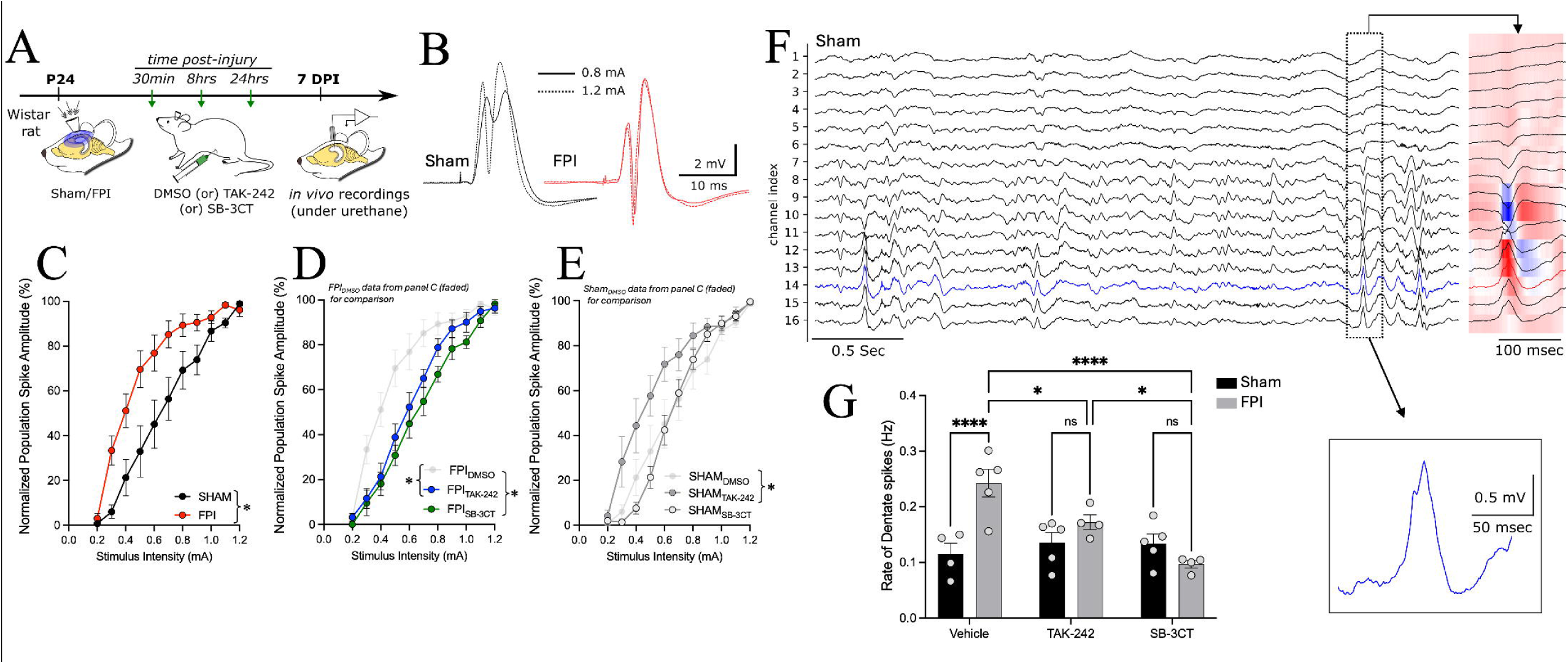
TLR4 regulation of MMP-3 activity enhances dentate network excitability after brain injury. A) Experimental timeline for *in vivo* experiments examining dentate excitability and throughput. B) Representative traces showing DG population spike from sham and injured animals in response to 800 µA stimulus (bold lines) and 1200 µA (dotted lines). C) Input-output curve illustrating early saturation of population spike amplitude in injured animals (red) compared to sham controls. D) Inhibition of TLR4 or MMP-9 effectively mitigates these changes. E) MMP-9 inhibition (light grey) did not affect input-output curves in sham animals, TLR4 inhibition increased network excitability of sham animals (grey). Sham and FPI data from panel C are included as faded traces in D (FPI) and E (Sham) for comparison. F) Representative trace from 16 channel silicon probe recording local field potentials from cortex to hippocampus. Current-source density plot shows the sink and source of an individual dentate spike. inset-a single dentate spike recorded in the GC layer. G) Quantification of dentate spikes shows a significant increase in frequency in injured animals. Inhibition of TLR4 or MMP-9 effectively reduced changes in dentate spike frequency in injured animals. n = 5-6 rats/group (A-E) and 4-5 rats/ group (F-G). * indicates<0.05, **** indicates *P*<0.0001 by TW-ANOVA with Šídáks multiple comparisons post hoc tests. Data points in panels G represent individual animals.

Enhanced network excitability combined with deficits in synaptic inhibition can increase dentate throughput. We assessed DG throughput by examining dentate spikes, a distinct DG population event driven by entorhinal cortex [53,54]. Linear 16-channel silicon probe recordings of dentate spikes in anesthetized rats (Figure 4F-G) revealed a significant increase in the frequency of dentate spikes one-week after FPI (in Hz: Sham: 0.11±0.019; FPI 0.24±0.025, n = 4-5 rats/group, *P=* 0.006 by two tailed unpaired t-test), identifying enhanced dentate throughput following FPI. To test if inhibition of either TLR4 signaling or MMP-9 activity early after injury would reduce changes to DG throughput, we examined dentate spikes in TAK-242 or SB-3CT treated sham/injured rats. Our experiments demonstrate that early inhibition of either TLR4 signaling or MMP-9 activity effectively prevented the post injury increase in dentate spike frequency without impacting it in sham controls (n = 4-5 rats/group; Effect of Interaction F (2,21) = 9.975, *P*<0.001 by Two-way ANOVA, Figure 4G).

We previously demonstrated that early post-injury increase in DGC calcium permeable AMPA currents and deficits in working memory were mitigated by TLR4 inhibition [13]. To assess whether the TLR4-MMP9 signaling impacts synaptic plasticity, we examined longterm potentiation (LTP) at perforant path-DGC synapses in anesthetized rats *in vivo* using theta-burst stimulation (TBS) [38]. In the first 10 minutes after TBS, the field EPSP slope normalized to the maximum slope at baseline trended lower but was not statistically different between sham and FPI. However, there was a reduction in the field EPSP slope at 60 min, 120 min, and 180 min post-TBS indicating impairment in LTP maintenance (n = 6 rats/group, Effect of injury: F (1,10) = 32.79, *P*=0.0002 by Two-way RM ANOVA with Šídák’s multiple comparisons 10 min: *P* =0.065, 60 min: *P* = 0.003, 120 min: *P* = 0.002, 180 min: *P* = 0.125, Figure 5A-E). Crucially, systemic antagonism of either TLR4 or MMP-9 early after FPI effectively prevented deficits in LTP in injured rats, demonstrating contribution of TLR4-MMP-9 axis to LTP deficits after brain injury (Effect of treatment: F(2,13) = 18, *P*=0.0002; *n*= 5-6 rats/group; See Supplementary table 4, Figure 5A-E). Consistent with the paradoxical effects of TAK-242 on sEPSC and network excitability in sham controls, TAK-242 treatment resulted in deficits in LTP in sham rats (Effect of treatment: F(2,13) = 8.046, *P*=0.0053; *n*= 5-6 rats/group; Figure 5E). Once again, unlike TAK242, SB-3CT treatment failed to impact LTP Sham rats, paralleling on the differential effects of the treatments on sEPSC frequency, population spike amplitude and dentate spike frequency in sham animals. These findings demonstrate that TLR4-MMP-9 dependent post-traumatic circuit changes compromise DG glutamatergic circuit function and plasticity.

**Figure 5:**
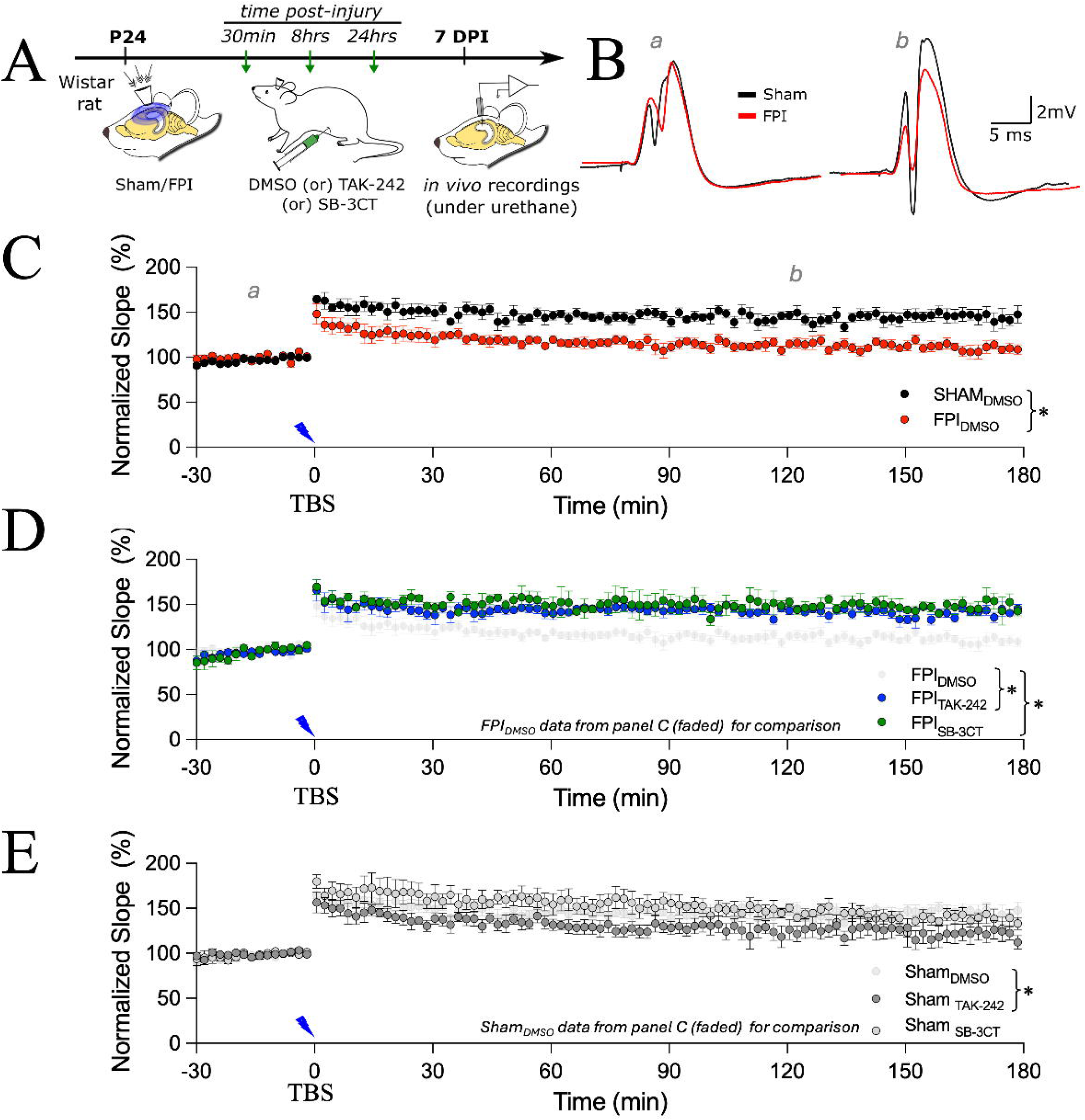
Posttraumatic deficits in long-term potentiation are improved by early inhibition of TLR4 or MMP-3. A) Experimental timeline for *in vivo* assessment of long-term potentiation in perforant path synapses to DGCs. B) Representative traces from Sham (black) and FPI rats (red) at 10 minutes before (a) and 120 minutes after (b) LTP induction. C) Injured rats exhibit a significant decrease in fEPSP slope values after 60 minutes(red) compared to sham controls (black). D) Inhibition of TLR4 (blue) or MMP-9 (green) significantly improved LTP in injured animals. E) MMP-9 inhibition did not affect LTP in SHAM controls. TLR4 inhibition reduced LTP at 60 min and 180 min after induction. Note that Sham and FPI data from panel C are included as faded traces in D (FPI) and E (Sham) for comparison. n = 5-6 rats/group * indicates *P* <0.05 by TW-RM ANOVA.

### Targeting TLR4 or MMP-9 axis early after brain injury limits long-term spatial memory impairments

Since we identify post injury increase in dentate spikes which can compromise hippocampal information processing [55–58], and blocking TLR4 and MMP-9 early after injury prevent this change, we sought to determine if TLR4 and MMP-9 antagonists could alleviate long term cognitive deficits after brain injury. We adopted a Barnes maze spatial learning task which has been shown to rely on intact DG function[59,60]. One month after sham/FPI, brain injured rats exhibited a longer latency to find the escape hole and traveled a greater total distance compared to sham controls early during training but reached control levels of task performance by Day 3 (See Supplementary table 5, *n* =10 rats / group, *P*<0.05 by TW-RM ANOVA, Figure 6A-D). Additionally, there was no difference in task performance during trials on Day 4 and Day 7, indicating that task retention is not impaired (Supplementary figure 6B-C). Although there were no differences in speed of locomotion (Supplementary figure 6D), injured rats showed increased thigmotaxis on Days 1 and 2 (Time spent in thigmotaxis zone in seconds: Sham_DMSO_ – 24.73±5.42, FPI_DMSO_ – 64.99±16.96; *n*= 10 rats/group, Effect of injury–F (1,18) = 6.841, *P*<0.05 by Two-way-RM ANOVA, Figure 6I). Since injured rats showed a significant increase in both thigmotaxis and latency to find the escape on Day 2 of task acquisition, we examined the use of spatial strategy for navigation using a 11-point scoring system [42]. Analysis of search strategies on Day 2 showed that injured animals employed random or serial strategies more often, demonstrating specific deficits in ability to recall spatial cues for navigation (Search strategy score_Day2: Sham_DMSO_ – 7.7±0.73, FPI_DMSO_ – 2.4±0.47; *n*= 10 rats/group, Effect of injury– F (1,18) = 17, *P*<0.01 by Two-way RM ANOVA, Figure.6E,H). These results suggest a mild delay in task acquisition after brain injury which is apparent on Day 2 of the task.

**Figure 6:**
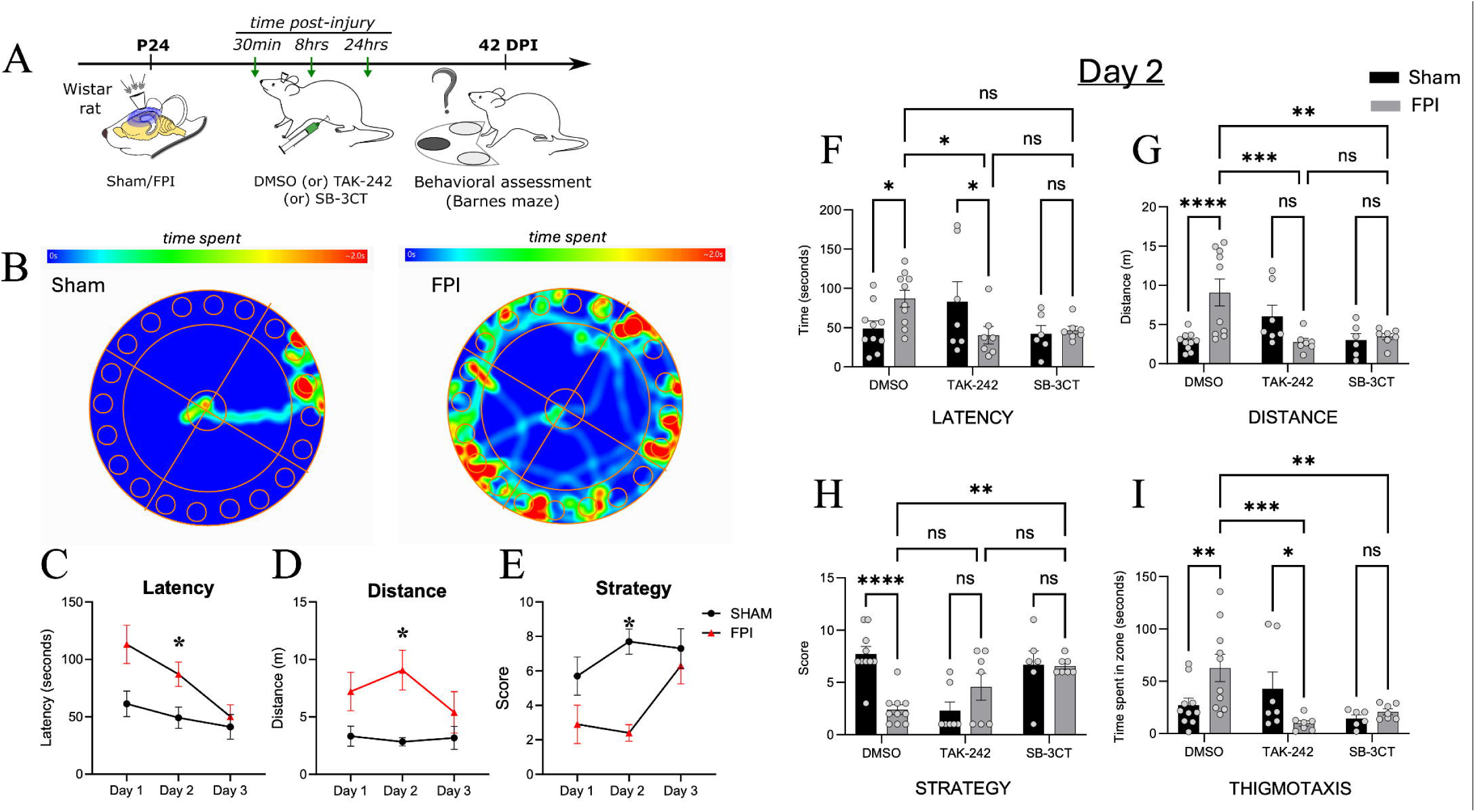
Early inhibition of TLR4 or MMP-3 improves spatial learning after brain injury. A) Experimental timeline for testing spatial learning using Barnes maze. B) Representative heat plot showing rats trajectory during the task on day 2. C-E) Injured animals showed deficits in latency to escape, distance travelled, and search strategy on day 2. * indicates P<0.05 by RM-ANOVA. F-H) Summary plots of effect of TLR4 or MMP-9 inhibition on latency to escape (F), distance travelled (G), and search strategy (H) on day 2. I) Injured animals showed increased thigmotaxis on day 2 which was attenuated with TLR4 or MMP-9 inhibition. n = 6-10 rats/group. * indicates P<0.05, ** indicates *P*<0.01, *** indicates *P*<0.001, **** indicates *P*<0.0001 by TW-ANOVA with Šídáks multiple comparisons post hoc tests. Data points in panels F-I represent individual animals.

We focused our analysis on Day 2 of task acquisition when the injury-induced impairment is most apparent. Early inhibition of TLR4 signaling or MMP-9 activity prevented post-injury increases in latency to escape, and distance traveled observed on Day 2 (n = 6-10 rats/group; *P*<0.05 by Two-way RM ANOVA, Figure 6F-G, Supplementary table 3 and Supplementary figure 6A). While the trend in ability of TLR4 inhibition to improve the strategy score in injured animals did not reach statistical significance (Figure 6H), the search strategy scores in injured animals significantly improved with MMP-9 inhibition (Figure 6H). Importantly, while MMP-9 did not have any adverse effects on sham controls, TLR4 inhibition showed a trend towards increased latency, distance travelled and thigmotaxis, and a lower spatial search strategy score (Figure 6F-I) which did not reach statistical significance. Collectively, these findings demonstrate that early inhibition of the TLR4-MMP-9 signaling axis effectively mitigates mild long term spatial learning deficits observed after FPI.

## DISCUSSION

Understanding how immune signaling modulates neuronal function after brain injury is critical for developing effective therapeutic strategies to minimize the long-term sequelae of TBI. Aside from the widespread secondary injury driven by glial mediators of neuroinflammation [1,2], we previously showed that the neuronal expression of innate immune receptor TLR4 is upregulated in the hippocampus early after injury, contributing to long-term seizure susceptibility and impaired working memory [12–14]. Here, we provide mechanistic evidence that TLR4 signaling mediates rapid, circuit-level changes in both excitatory and inhibitory synaptic transmission after injury. Using a combination of pharmacology in rats and mouse genetics we demonstrate a central role for TLR4 expressed in CaMKII-positive excitatory neurons in mediating the posttraumatic circuit changes through the upregulation of MMP-9. Importantly, we show that either TLR4 or MMP-9 inhibition early after brain injury not only reduces these pathological changes one-week after injury, but improves deficits in spatial learning 1-month after injury.

### TLR4 Signaling Augments Glutamatergic inputs to DGCs after brain injury

TLR4 signaling is well established as a driver of neuronal excitability [3,61–65]. Our prior works showed that TLR4 signaling enhances AMPA currents in granule cells contributing to DG network hyperexcitability [12,13]. Examining circuit-level changes in animals one week after injury, this study identifies an aberrant increase in excitatory inputs to DGCs in the dorsal hippocampus. Considering that perforant path fibers are severed in coronal hippocampal slice preparations [66], the post-injury increase in spontaneous EPSCs and action potential-independent miniature events observed here indicate an increase in synaptic contacts, consistent with early axonal sprouting [46], or elevated mossy cell excitability previously reported after FPI [67]. Our findings highlight a critical role of TLR4 signaling in this process. First, we show that early pharmacological inhibition of TLR4 mitigates changes in both spontaneous and miniature EPSC frequency. We further demonstrate that transient activation of TLR4 signaling in acute brain slices using LPS-B5 is sufficient to augment sEPSCs frequency to levels comparable to those seen in injured animals. Furthermore, by leveraging TLR4^fl/fl^: CaMKIIa-CreERT2 mice, we demonstrate that selective deletion of TLR4 in glutamatergic neurons alone is sufficient to mitigate the post-injury increase in sEPSC frequency. Together, these findings show that neuronal TLR4 is necessary to enhance glutamatergic drive to DGCs after brain injury.

### A Novel Neuro-Immune-enzyme Signaling Cascade Underlies Early Post-Traumatic Glutamatergic Circuit Plasticity

To elucidate the mechanism by which TLR4 enhances glutamatergic drive to DGCs after injury, we focused on MMP-9 as a potential downstream effector. MMP-9 co-localizes with excitatory synapses in an activity-dependent manner, and increased MMP-9 activity is known to dysregulate glutamatergic synaptic transmission [40,52,68–70]. We previously reported that neuronal TLR4 enhances calcium-permeable AMPA currents in DGCs post-injury by increasing the surface expression of GluA1 subunits [13]. Notably, MMP-9 activity has been shown to regulate AMPA receptor mobility and synaptic accumulation of GluA1 subunits following chemically induced LTP [71]. In addition, TLR4 ligands such as high-mobility group box 1 (HMGB1) and lipopolysaccharide (LPS) are known to enhance MMP-9 expression in cultures and following ischemic insult [15,16] making MMP9 an attractive candidate effector of the synaptic effects of TLR4.

Here, we show that pharmacologically inhibiting TLR4 signaling (rats) or deleting TLR4 in excitatory neurons (mice) both limit post-injury increases in MMP-9 activity identifying that TLR4 signaling is necessary for injury-induced increase in MMP9. Furthermore, using LPS-B5 to selectively activate TLR4, we demonstrate that activation of TLR4 is sufficient to increase MMP-9 mRNA levels in the dentate gyrus of naïve rats in an NF-κB dependent manner. Thus, as summarized in Figure 7, our findings identify a novel TLR4/NF-κB-dependent transcriptional pathway that upregulates MMP-9 activity and promotes glutamatergic circuit reorganization following brain injury. Consistent with this, inhibition of MMP-9 with SB-3CT effectively reduced most of the glutamatergic circuit changes in the DG resulting from direct activation of TLR4 signaling in slices (LPS-B5) and brain injury *in vivo*.

**Figure 7:**
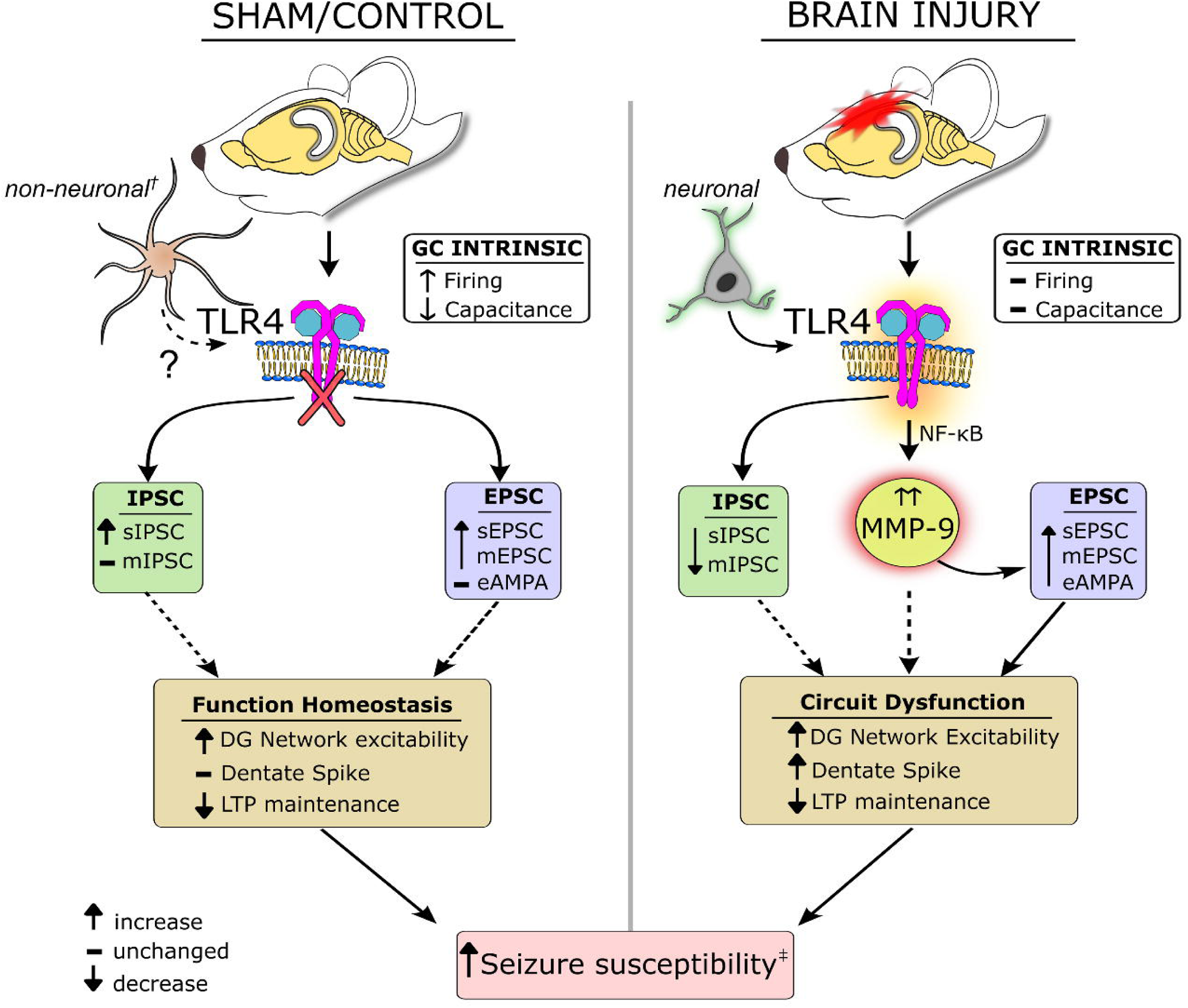
Graphical Summary of TLR4 regulation of dentate gyrus function in control and injured brain. In uninjured brain, pharmacological inhibition of TLR4 signaling increases the frequency of both excitatory and inhibitory currents and enhances network excitability and impairs longterm potentiation without altering MMP-9 activity. Genetic deletion of neuronal TLR4 alters intrinsic properties without impacting synaptic physiology. Together these data are consistent with non-neuronal TLR4 signaling contributing to basal synaptic homeostasis in line with prior ex vivo studies [14]. In the injured brain, augmented neuronal TLR4 signaling acts through MMP-9 to enhance glutamatergic signaling, network excitability and dentate spikes and impairs LTP maintenance. Mechanistically, inhibitory changes are driven by TLR4 in an MMP-9 independent manner. Together, these findings demonstrate that TLR4 signaling regulates DG function and seizure susceptibility [13] through distinct, context-dependent cellular mechanisms in the healthy and injured brain.† refers to [14] ‡ refers to [13].

### Mechanistic Divergence in TLR4 regulation of glutamatergic and GABAergic circuits

In contrast to the glutamatergic inputs, we observed a decrease in the frequency of inhibitory inputs to dorsal DGCs one week after FPI. This stands in direct contrast to our previous findings from the ventral hippocampus [36,45,46], highlighting potential regional differences in functional plasticity between dorsal and ventral hippocampal inhibitory circuits [72]. While TLR4 inhibition reduced post-traumatic changes in IPSCs, this effect was independent of MMP-9 activity. Although enhances MMP-9 activity is implicated in hypofunction of parvalbumin expressing inhibitory neurons and perineuronal net remodeling in neurodevelopmental disorders [73], we find that blocking MMP9 does not impact changes in inhibitory synaptic inputs after brain injury. Notably, our findings indicate a divergence of TLR4-dependent mechanisms for glutamatergic and GABAergic circuit reorganization, with MMP-9-independent mechanisms contributing to changes in inhibitory synapses (Figure 7). Interestingly, unlike after brain injury, selective activation of TLR4 with LPS-B5 in slices from sham rats did not alter sIPSC frequency in DGCs raising the possibility that injury-driven loss of hilar interneurons contributes to the reduction in sIPSC after brain injury [7,47]. However, mIPSC frequency was increased following LPS-B5 incubation suggesting perturbations in release probability or presynaptic calcium [44,74–76]. Curiously, mice with TLR4 deletion developed posttraumatic depolarization of RMP not observed in cre^-ve^ controls suggesting TLR4 signaling may contribute to cell intrinsic compensatory processes. Future studies should determine whether neuronal or glial TLR4 signaling mediates the TLR4 dependent post-injury changes in inhibition and how this impacts dentate function and behaviors.

### Network Consequences and behavioral impact of TLR4/MMP-9 pathway

The dentate gyrus is a central node in hippocampal memory processing, regulating the flow of information through the hippocampal circuit [56,77,78]. Consequently, perturbations in DG circuit function can directly impact cognitive processes. Our earlier studies showed enhanced perforant path evoked EPSC amplitude in DGCs after FPI, suggesting stronger entorhinal-dentate coupling in the injured brain, coupled with deficits in working memory [12–14]. Here we advance these findings using in vivo approaches to identify increased dentate network excitability early after injury. We further show that while LTP induction was unaffected after injury, LTP maintenance was compromised in injured rats. Indeed the lack of deficit in LTP induction argues against a role for synaptic saturation in mediating LTP deficits. Rather, we find selective deficits in LTP maintenance after brain injury indicating altered transcriptional processes. Importantly, we report an increase in dentate spike frequency early after brain injury indicating abnormal entorhinal-dentate throughput. Since dentate spikes are critical in hippocampal information processing, memory formation, and consolidation [53–57,79,80], abnormal dentate spikes could impair memory processing after brain injury. Supporting this proposal, treatments that reduced dentate spike frequency in injured animals also improved spatial learning deficits. Crucially, dentate spikes are also increased in experimental models of epilepsy [81], suggesting that these events may serve as biomarkers for post-traumatic epileptogenesis alongside the pathological high frequency oscillations and altered sleep spindles identified previously [82,83]. In summary, inhibition of the novel TLR4-MMP9 signaling axis after brain injury preserves dentate circuit excitability and long-term potentiation while rescuing spatial memory deficits (Figure 7). These results highlight the manipulation of this pathway as a promising therapeutic strategy for post-traumatic recovery.

### A Novel Constitutive Role for TLR4 Signaling in Basal Circuit Homeostasis

Our study reveals a fundamental role for basal TLR4 signaling in maintaining dentate circuit homeostasis. We found that inhibition of TLR4 signaling in uninjured animals consistently altered synaptic inputs, network excitability, and plasticity. Specifically, timed and selective deletion of TLR4 in glutamatergic neurons in a mature circuit caused a genotype-dependent reduction in granule cell capacitance and increased firing rates (Figure 7). This indicates a constitutive role for neuronal TLR4 in limiting basal excitability. A reduction in capacitance with increased firing in the absence of changes in action potential threshold parallels observations in parvalbumin neurons in epilepsy models, where regulation of perineuronal nets alters neuronal firing through effects on cell capacitance[41,84,85].

Indeed, our prior studies have found that ability of TLR4 blockers to increase dentate excitability in uninjured controls is abolished when glial metabolic processes are disrupted[14]. Our finding that TLR4 or MMP9 antagonism does not impact dentate spike frequency in uninjured controls, is consistent with lack of TLR4 modulation of perforant path evoked AMPA currents in the absence of injury. Nevertheless, the basal TLR4 regulation of network homeostasis appears critical as blocking TLR4 enhances network excitability, impairs LTP maintenance and spatial memory (Figure 7). The potential for TLR4 inhibitors to limit the progression of epilepsy and Alzheimer’s disease [11,61,86,87], highlights the need to understand non-canonical TLR4 signaling in the healthy brain. Future research should focus on the underlying mechanisms of basal TLR4-mediated neuronal regulation.

Collectively, our results position the TLR4-NFkB-MMP-9 axis as a central driver of TBI-induced circuit dysfunction, providing a targetable molecular pathway to serve as a therapeutic target for mitigating post-traumatic epilepsy and associated cognitive deficits.

## Declarations

### Ethics approval and consent to participate

All experiments were performed in accordance with IACUC protocols approved by the University of California-Riverside, CA (Approval number #20210005/35) and in keeping with the ARRIVE guidelines.

### Consent for publication

Not applicable.

### Availability of data and materials

All data are available from the corresponding author upon reasonable request.

### Authors’ contributions

D.S., E.C., and L.D., performed experiments; D.S., E.C., L.D., E.G., Y.L., and R.J., analyzed data; D.S., I.E., and V.S., interpreted results of experiments; D.S. prepared figures; D.S., V.S. conception and design of research; D.S. and V.S., drafted manuscript; D.S., I.E., and V.S. finalized and approved manuscript.

## Acknowledgements

We thank Dr. Leszek Kaczmarek (Nencki Institute, Warsaw, Poland) for his expert advice on *in situ* zymography. We also thank Dr. Amanda Schott and Dr. Jordan S Farrell (Harvard Medical School) for their expertise and help with Dentate spike detection algorithms.

## List of Abbreviations

aCSF: Artificial Cerebrospinal Fluid
AMPAR: α-Amino-3-hydroxy-5-methyl-4-isoxazolepropionic Acid Receptor
ANOVA: Analysis of Variance
CA1 SR: CA1 Stratum Radiatum
CaMKIIa: Calcium/Calmodulin-Dependent Protein Kinase II Alpha
CreERT2: Cre Recombinase Estrogen Receptor T2
DG: Dentate Gyrus
DQ Gelatinase: Dye Quenched fluorogenic substrate for zymography
ECM: Extracellular Matrix
EPSC: Excitatory Postsynaptic Current
FPI: Fluid Percussion Injury
HMGB1: High-Mobility Group Box 1
IEI: Inter-Event Interval
IPSC: Inhibitory Postsynaptic Current
JSH-23: NF-κB Inhibitor
K-S test: Kolmogorov-Smirnov Test
LPS: Lipopolysaccharide
LTP: Long Term Potentiation
MMP-9: Matrix Metalloproteinase-9
NF-κB: Nuclear Factor Kappa B
TAK-242: TLR4 Antagonist
TBS: Theta Burst Stimulation
TBI: Traumatic Brain Injury
TTX: Tetrodotoxin

## REFERENCES

1. Sharma R, Leung WL, Zamani A, O’brien TJ, Espinosa PMC, Semple BD. Neuroinflammation in post-traumatic epilepsy: Pathophysiology and tractable therapeutic targets. Brain Sciences. 2019 Nov 1;9(11). doi:10.3390/brainsci9110318

2. Mukherjee S, Arisi GM, Mims K, Hollingsworth G, O’Neil K, Shapiro LA. Neuroinflammatory mechanisms of post-traumatic epilepsy. Journal of Neuroinflammation. 2020 Jun 17;17(1). doi:10.1186/s12974-020-01854-w PubMed PMID: 32552898.

3. Webster KM, Sun M, Crack P, O’Brien TJ, Shultz SR, Semple BD. Inflammation in epileptogenesis after traumatic brain injury. Journal of Neuroinflammation. 2017 Jan 13;14(1). doi:10.1186/s12974-016-0786-1 PubMed PMID: 28086980.

4. Simon DW, McGeachy MJ, Baylr H, Clark RSB, Loane DJ, Kochanek PM. The far-reaching scope of neuroinflammation after traumatic brain injury. Nature Reviews Neurology. 2017 Mar 1;13(3):171–91. doi:10.1038/nrneurol.2017.13 PubMed PMID: 28186177.

5. Lowenstein D, Thomas M, Smith D, McIntosh T. Selective vulnerability of dentate hilar neurons following traumatic brain injury: a potential mechanistic link between head trauma and disorders of the hippocampus. J Neurosci. 1992 Dec 1;12(12):4846–53. doi:10.1523/JNEUROSCI.12-12-04846.1992

6. McIntosh, Saatman, Raghupathi, Graham, Smith, Lee, et al. The Dorothy Russell Memorial Lecture* The molecular and cellular sequelae of experimental traumatic brain injury: pathogenetic mechanisms. Neuropathology Appl Neurobio. 1998 Aug;24(4):251–67. doi:10.1046/j.1365-2990.1998.00121.x

7. Toth Z, Hollrigel GS, Gorcs T, Soltesz I. Instantaneous Perturbation of Dentate Interneuronal Networks by a Pressure Wave-Transient Delivered to the Neocortex. 1997.

8. Atkins CM. Decoding Hippocampal Signaling Deficits After Traumatic Brain Injury. Transl Stroke Res. 2011 Dec;2(4):546–55. doi:10.1007/s12975-011-0123-z

9. Hunt RF, Boychuk JA, Smith BN. Neural circuit mechanisms of post–traumatic epilepsy. Front Cell Neurosci. 2013;7. doi:10.3389/fncel.2013.00089

10. Paterno R, Folweiler KA, Cohen AS. Pathophysiology and Treatment of Memory Dysfunction After Traumatic Brain Injury. Curr Neurol Neurosci Rep. 2017 Jul;17(7):52. doi:10.1007/s11910-017-0762-x

11. Neuberger EJ, Gupta A, Subramanian D, Korgaonkar AA, Santhakumar V. Converging early responses to brain injury pave the road to epileptogenesis. Journal of Neuroscience Research. 2019 Nov 1;97(11):1335–44. doi:10.1002/jnr.24202 PubMed PMID: 29193309.

12. Li Y, Korgaonkar AA, Swietek B, Wang J, Elgammal FS, Elkabes S, et al. Toll-like receptor 4 enhancement of non-NMDA synaptic currents increases dentate excitability after brain injury. Neurobiology of Disease. 2015 Feb 1;74:240–53. doi:10.1016/j.nbd.2014.11.021 PubMed PMID: 25497689.

13. Korgaonkar AA, Li Y, Sekhar D, Subramanian D, Guevarra J, Swietek B, et al. Toll-like Receptor 4 Signaling in Neurons Enhances Calcium-Permeable α-Amino-3-Hydroxy-5-Methyl-4-Isoxazolepropionic Acid Receptor Currents and Drives Post-Traumatic Epileptogenesis. Annals of Neurology. 2020 Apr 1;87(4):497–515. doi:10.1002/ana.25698 PubMed PMID: 32031699.

14. Korgaonkar AA, Nguyen S, Li Y, Sekhar D, Subramanian D, Guevarra J, et al. Distinct cellular mediators drive the Janus faces of toll-like receptor 4 regulation of network excitability which impacts working memory performance after brain injury. Brain, behavior, and immunity. 2020 Aug;88:381–95. doi:10.1016/j.bbi.2020.03.035 PubMed PMID: 32259563.

15. Yang CC, Lin CC, Hsiao LD, Kuo JM, Tseng HC, Yang CM. Lipopolysaccharide-induced matrix metalloproteinase-9 expression associated with cell migration in rat brain astrocytes. International Journal of Molecular Sciences. 2020 Jan 1;21(1). doi:10.3390/ijms21010259 PubMed PMID: 31905967.

16. Qiu J, Xu J, Zheng Y, Wei Y, Zhu X, Lo EH, et al. High-mobility group box 1 promotes metalloproteinase-9 upregulation through toll-like receptor 4 after cerebral ischemia. Stroke. 2010 Sep;41(9):2077–82. doi:10.1161/STROKEAHA.110.590463 PubMed PMID: 20671243.

17. Huntley GW. Synaptic circuit remodelling by matrix metalloproteinases in health and disease. Nature Reviews Neuroscience. 2012 Nov;13(11):743–57. doi:10.1038/nrn3320 PubMed PMID: 23047773.

18. Ethell IM, Ethell DW. Matrix metalloproteinases in brain development and remodeling: synaptic functions and targets. Journal of neuroscience research. 2007 Oct;85(13):2813–23. doi:10.1002/jnr.21273 PubMed PMID: 17387691.

19. Szklarczyk A, Lapinska J, Rylski M, Mckay RDG, Kaczmarek L. Matrix Metalloproteinase-9 Undergoes Expression and Activation during Dendritic Remodeling in Adult Hippocampus. 2002.

20. Dityatev A, Schachner M. Extracellular matrix molecules and synaptic plasticity. Nature reviews Neuroscience. 2003 Jun;4(6):456–68. doi:10.1038/nrn1115 PubMed PMID: 12778118.

21. Minta K, Brinkmalm G, Al Nimer F, Thelin EP, Piehl F, Tullberg M, et al. Dynamics of cerebrospinal fluid levels of matrix metalloproteinases in human traumatic brain injury. Sci Rep. 2020 Oct 22;10(1):18075. doi:10.1038/s41598-020-75233-z

22. Wang X, Jung J, Asahi M, Chwang W, Russo L, Moskowitz MA, et al. Effects of Matrix Metalloproteinase-9 Gene Knock-Out on Morphological and Motor Outcomes after Traumatic Brain Injury. 2000.

23. Hayashi T, Kaneko Y, Yu S, Bae E, Stahl CE, Kawase T, et al. Quantitative analyses of matrix metalloproteinase activity after traumatic brain injury in adult rats. Brain Research. 2009 Jul;1280:172–7. doi:10.1016/j.brainres.2009.05.040

24. Grossetete M, Phelps J, Arko L, Yonas H, Rosenberg GA. ELEVATION OF MATRIX METALLOPROTEINASES 3 AND 9 IN CEREBROSPINAL FLUID AND BLOOD IN PATIENTS WITH SEVERE TRAUMATIC BRAIN INJURY. Neurosurgery. 2009 Oct;65(4):702–8. doi:10.1227/01.NEU.0000351768.11363.48

25. Reinhard SM, Razak K, Ethell IM. A delicate balance: Role of MMP-9 in brain development and pathophysiology of neurodevelopmental disorders. Frontiers in Cellular Neuroscience. 2015 Jul 29;9(JULY). doi:10.3389/fncel.2015.00280

26. Vafadari B, Salamian A, Kaczmarek L. MMP-9 in translation: from molecule to brain physiology, pathology, and therapy. Journal of Neurochemistry. 2016 Oct 1;139:91–114. doi:10.1111/jnc.13415 PubMed PMID: 26525923.

27. Pijet B, Stefaniuk M, Kostrzewska-Ksiezyk A, Tsilibary PE, Tzinia A, Kaczmarek L. Elevation of MMP-9 Levels Promotes Epileptogenesis After Traumatic Brain Injury. Molecular Neurobiology. 2018 Dec 1;55(12):9294–306. doi:10.1007/s12035-018-1061-5 PubMed PMID: 29667129.

28. Wilczynski GM, Konopacki FA, Wilczek E, Lasiecka Z, Gorlewicz A, Michaluk P, et al. Important role of matrix metalloproteinase 9 in epileptogenesis. Journal of Cell Biology. 2008 Mar 10;180(5):1021–35. doi:10.1083/jcb.200708213 PubMed PMID: 18332222.

29. Dityatev A, Fellin T. Extracellular matrix in plasticity and epileptogenesis. Neuron Glia Biol. 2008 Aug;4(3):235–47. doi:10.1017/S1740925X09000118

30. Jia F, Pan Y Hua, Mao Q, Liang Y Min, Jiang J Yao. Matrix metalloproteinase-9 expression and protein levels after fluid percussion injury in rats: The effect of injury severity and brain temperature. Journal of neurotrauma. 2010 Jun;27(6):1059–68. doi:10.1089/neu.2009.1067 PubMed PMID: 20233042.

31. Corrubia L, Huang A, Nguyen S, Shiflett MW, Jones MV, Ewell LA, et al. Early deficits in dentate circuit and behavioral pattern separation after concussive brain injury. Experimental Neurology. 2023 Dec;370:114578. doi:10.1016/j.expneurol.2023.114578

32. Witgen BM, Lifshitz J, Smith ML, Schwarzbach E, Liang SL, Grady MS, et al. Regional hippocampal alteration associated with cognitive deficit following experimental brain injury: A systems, network and cellular evaluation. Neuroscience. 2005;133(1):1–15. doi:10.1016/j.neuroscience.2005.01.052

33. Jia F, Yin YH, Gao GY, Wang Y, Cen L, Jiang JY. MMP-9 inhibitor SB-3CT attenuates behavioral impairments and hippocampal loss after traumatic brain injury in rat. Journal of Neurotrauma. 2014 Jul 1;31(13):1225–34. doi:10.1089/neu.2013.3230 PubMed PMID: 24661104.

34. Fumihiro J, Masaaki K, Toshiyuki T, Yoshihiko T, Takahiro K, Satoru A. Investigation of the unique metabolic fate of ethyl (6R)-6-[N-(2-chloro-4-fluorophenyl)sulfamoyl]cyclohex-1-ene-1-carboxylate (TAK-242) in rats and dogs using two types of 14C-labeled compounds having different labeled positions. Arzneimittelforschung. 2011 Nov 27;61(08):458–71. doi:10.1055/s-0031-1296228

35. Hadass O, Tomlinson BN, Gooyit M, Chen S, Purdy JJ, Walker JM, et al. Selective Inhibition of Matrix Metalloproteinase-9 Attenuates Secondary Damage Resulting from Severe Traumatic Brain Injury. PLoS ONE. 2013 Oct 23;8(10). doi:10.1371/journal.pone.0076904 PubMed PMID: 24194849.

36. Gupta A, Elgammal FS, Proddutur A, Shah S, Santhakumar V. Decrease in Tonic Inhibition Contributes to Increase in Dentate Semilunar Granule Cell Excitability after Brain Injury. J Neurosci. 2012 Feb 15;32(7):2523–37. doi:10.1523/JNEUROSCI.4141-11.2012

37. Hirschfeld M, Ma Y, Weis JH, Vogel SN, Weis JJ. Cutting edge: repurification of lipopolysaccharide eliminates signaling through both human and murine toll-like receptor 2. J Immunol. 2000 Jul 15;165(2):618–22. doi:10.4049/jimmunol.165.2.618 PubMed PMID: 10878331.

38. Cooke SF, Wu J, Plattner F, Errington M, Rowan M, Peters M, et al. Autophosphorylation of αCaMKII is not a general requirement for NMDA receptor-dependent LTP in the adult mouse. Journal of Physiology. 2006 Aug;574(3):805–18. doi:10.1113/jphysiol.2006.111559 PubMed PMID: 16728448.

39. Schott AL, Esfahany KN, Grocott JM, Farrell JS. Toothy: an interactive platform for dentate spike curation. bioRxiv. 2025 Oct 3;2025.10.03.680300. doi:10.1101/2025.10.03.680300 PubMed PMID: 41256680; PubMed Central PMCID: PMC12621741.

40. Gawlak M, Górkiewicz T, Gorlewicz A, Konopacki FA, Kaczmarek L, Wilczynski GM. High resolution in situ zymography reveals matrix metalloproteinase activity at glutamatergic synapses. Neuroscience. 2009 Jan 12;158(1):167–76. doi:10.1016/j.neuroscience.2008.05.045 PubMed PMID: 18588950.

41. Proddutur A, Nguyen S, Yeh CW, Gupta A, Santhakumar V. Reclusive chandeliers: Functional isolation of dentate axo-axonic cells after experimental status epilepticus. Progress in Neurobiology. 2023 Dec;231:102542. doi:10.1016/j.pneurobio.2023.102542

42. Rodríguez Peris L, Scheuber MI, Shan H, Braun M, Schwab ME. Barnes maze test for spatial memory: A new, sensitive scoring system for mouse search strategies. Behavioural Brain Research. 2024 Feb;458:114730. doi:10.1016/j.bbr.2023.114730

43. Harrison FE, Reiserer RS, Tomarken AJ, McDonald MP. Spatial and nonspatial escape strategies in the Barnes maze. Learn Mem. 2006 Nov;13(6):809–19. doi:10.1101/lm.334306

44. Hunt RF, Scheff SW, Smith BN. Synaptic Reorganization of Inhibitory Hilar Interneuron Circuitry after Traumatic Brain Injury in Mice. J Neurosci. 2011 May 4;31(18):6880–90. doi:10.1523/JNEUROSCI.0032-11.2011

45. Gupta A, Dovek L, Proddutur A, Elgammal FS, Santhakumar V. Long-Term Effects of Moderate Concussive Brain Injury During Adolescence on Synaptic and Tonic GABA Currents in Dentate Granule Cells and Semilunar Granule Cells. Front Neurosci. 2022;16:800733. doi:10.3389/fnins.2022.800733 PubMed PMID: 35360164; PubMed Central PMCID: PMC8964009.

46. Santhakumar V, Ratzliff ADH, Jeng J, Toth Z, Soltesz I. Long-term hyperexcitability in the hippocampus after experimental head trauma. Annals of Neurology. 2001 Dec;50(6):708–17. doi:10.1002/ana.1230

47. Santhakumar V, Bender R, Frotscher M, Ross ST, Hollrigel GS, Toth Z, et al. Granule cell hyperexcitability in the early post-traumatic rat dentate gyrus: the ‘irritable mossy cell’ hypothesis. The Journal of Physiology. 2000 Apr;524(1):117–34. doi:10.1111/j.1469-7793.2000.00117.x

48. Michaluk P, Wawrzyniak M, Alot P, Szczot M, Wyrembek P, Mercik K, et al. Influence of matrix metalloproteinase MMP-9 on dendritic spine morphology. J Cell Sci. 2011 Oct 1;124(Pt 19):3369–80. doi:10.1242/jcs.090852 PubMed PMID: 21896646.

49. Wiera G, Szczot M, Wojtowicz T, Lebida K, Koza P, Mozrzymas JW. Impact of matrix metalloproteinase-9 overexpression on synaptic excitatory transmission and its plasticity in rat CA3-CA1 hippocampal pathway. J Physiol Pharmacol. 2015 Apr;66(2):309–15. PubMed PMID: 25903961.

50. Nagy V, Bozdagi O, Matynia A, Balcerzyk M, Okulski P, Dzwonek J, et al. Matrix metalloproteinase-9 is required for hippocampal late-phase long-term potentiation and memory. Journal of Neuroscience. 2006 Feb 15;26(7):1923–34. doi:10.1523/JNEUROSCI.4359-05.2006 PubMed PMID: 16481424.

51. Bozdagi O, Nagy V, Kwei KT, Huntley GW. In vivo roles for matrix metalloproteinase-9 in mature hippocampal synaptic physiology and plasticity. Journal of Neurophysiology. 2007 Jul;98(1):334–44. doi:10.1152/jn.00202.2007 PubMed PMID: 17493927.

52. Wiera G, Mozrzymas JW. Extracellular Metalloproteinases in the Plasticity of Excitatory and Inhibitory Synapses. Cells. 2021 Aug 11;10(8). doi:10.3390/cells10082055 PubMed PMID: 34440823.

53. Bragin A, JandC G, Nadasdy Z, Van Landeghem M, Buzsaki G. Dentate EEG Spikes and Associated Interneuronal Population Bursts in the Hippocampal Hilar Region of the Rat. JOURNALOF NEUROPHYSIOLOGY. 1995.

54. Penttonen M, Kamondi A, Sik A, Acsády L, Buzsáki G. Feed-forward and feed-back activation of the dentate gyrus in vivo during dentate spikes and sharp wave bursts. Hippocampus. 1997;7(4):437–50. doi:10.1002/(SICI)1098-1063(1997)7:4%3C437::AID-HIPO9%3E3.0.CO;2-F PubMed PMID: 9287083.

55. Lensu S, Waselius T, Penttonen M, Nokia MS. Dentate spikes and learning: disrupting hippocampal function during memory consolidation can improve pattern separation. J Neurophysiol. 2019;121:131–9. doi:10.1152/jn.00696.2018.-Hippocampal

56. Farrell JS, Soltesz I. Noncanonical circuits, states, and computations of the hippocampus. Science. 2025 Sep 11;389(6765):eadv4420. doi:10.1126/science.adv4420

57. Nokia MS, Gureviciene I, Waselius T, Tanila H, Penttonen M. Hippocampal electrical stimulation disrupts associative learning when targeted at dentate spikes. Journal of Physiology. 2017 Jul 15;595(14):4961–71. doi:10.1113/JP274023 PubMed PMID: 28426128.

58. Farrell JS, Hwaun E, Dudok B, Soltesz I. Neural and behavioural state switching during hippocampal dentate spikes. Nature. 2024 Apr 18;628(8008):590–5. doi:10.1038/s41586-024-07192-8

59. Bott JB, Muller MA, Jackson J, Aubert J, Cassel JC, Mathis C, et al. Spatial reference memory is associated with modulation of theta-gamma coupling in the dentate Gyrus. Cerebral Cortex. 2016 Sep 1;26(9):3744–53. doi:10.1093/cercor/bhv177 PubMed PMID: 26250776.

60. Dovek L, Ahmadi M, Marrero K, Zagha E, Santhakumar V. Cellular and circuit features distinguish dentate gyrus semilunar granule cells and granule cells activated during contextual memory formation [Internet]. 2025 Jul 17. doi:10.7554/eLife.101428.2

61. Maroso M, Balosso S, Ravizza T, Liu J, Aronica E, Iyer AM, et al. Toll-like receptor 4 and high-mobility group box-1 are involved in ictogenesis and can be targeted to reduce seizures. Nature Medicine. 2010 Apr;16(4):413–9. doi:10.1038/nm.2127 PubMed PMID: 20348922.

62. Kleen JK, Holmes GL. Taming TLR4 may ease seizures. Nature Medicine. 2010 Apr;16(4):369–70. doi:10.1038/nm0410-369 PubMed PMID: 20376038.

63. Vezzani A, Balosso S, Ravizza T. Neuroinflammatory pathways as treatment targets and biomarkers in epilepsy. Nat Rev Neurol. 2019 Aug;15(8):459–72. doi:10.1038/s41582-019-0217-x

64. Okun E, Griffioen KJ, Mattson MP. Toll-like receptor signaling in neural plasticity and disease. Trends in Neurosciences. 2011 May;34(5):269–81. doi:10.1016/j.tins.2011.02.005 PubMed PMID: 21419501.

65. Ahmad A, Crupi R, Campolo M, Genovese T, Esposito E, Cuzzocrea S. Absence of TLR4 Reduces Neurovascular Unit and Secondary Inflammatory Process after Traumatic Brain Injury in Mice. Dhandapani KM, editor. PLoS ONE. 2013 Mar 28;8(3):e57208. doi:10.1371/journal.pone.0057208

66. Xiong G, Metheny H, Johnson BN, Cohen AS. A comparison of different slicing planes in preservation of major hippocampal pathway fibers in the mouse. Frontiers in Neuroanatomy. 2017 Nov 21;11. doi:10.3389/fnana.2017.00107

67. Howard AL, Neu A, Morgan RJ, Echegoyen JC, Soltesz I. Opposing Modifications in Intrinsic Currents and Synaptic Inputs in Post-Traumatic Mossy Cells: Evidence for Single-Cell Homeostasis in a Hyperexcitable Network. Journal of Neurophysiology. 2007 Mar;97(3):2394–409. doi:10.1152/jn.00509.2006

68. Dziembowska M, Milek J, Janusz A, Rejmak E, Romanowska E, Gorkiewicz T, et al. Activity-Dependent Local Translation of Matrix Metalloproteinase-9. J Neurosci. 2012 Oct 17;32(42):14538–47. doi:10.1523/JNEUROSCI.6028-11.2012

69. Dziembowska M, Wlodarczyk J. MMP9: A novel function in synaptic plasticity. The International Journal of Biochemistry & Cell Biology. 2012 May;44(5):709–13. doi:10.1016/j.biocel.2012.01.023

70. Stefaniuk M, Beroun A, Lebitko T, Markina O, Leski S, Meyza K, et al. Matrix Metalloproteinase-9 and Synaptic Plasticity in the Central Amygdala in Control of Alcohol-Seeking Behavior. Biological Psychiatry. 2017 Jun;81(11):907–17. doi:10.1016/j.biopsych.2016.12.026

71. Szepesi Z, Hosy E, Ruszczycki B, Bijata M, Pyskaty M, Bikbaev A, et al. Synaptically released matrix metalloproteinase activity in control of structural plasticity and the cell surface distribution of GluA1-AMPA receptors. PLoS ONE. 2014 May 22;9(5). doi:10.1371/journal.pone.0098274 PubMed PMID: 24853857.

72. Fanselow EE, Kubota Y, Wehr M, Yavorska I. Somatostatin-Expressing Inhibitory Interneurons in Cortical Circuits. Frontiers in Neural Circuits | www.frontiersin.org. 2016;10:76. doi:10.3389/fncir.2016.00076

73. Pirbhoy PS, Rais M, Lovelace JW, Woodard W, Razak KA, Binder DK, et al. Acute pharmacological inhibition of matrix metalloproteinase-9 activity during development restores perineuronal net formation and normalizes auditory processing in Fmr1 KO mice. Journal of Neurochemistry. 2020 Dec;155(5):538–58. doi:10.1111/jnc.15037

74. Sun DA, Deshpande LS, Sombati S, Baranova A, Wilson MS, Hamm RJ, et al. Traumatic brain injury causes a long-lasting calcium (Ca2+)-plateau of elevated intracellular Ca levels and altered Ca2+homeostatic mechanisms in hippocampal neurons surviving brain injury. Eur J of Neuroscience. 2008 Apr;27(7):1659–72. doi:10.1111/j.1460-9568.2008.06156.x

75. Weber JT. Altered Calcium Signaling Following Traumatic Brain Injury. Front Pharmacol. 2012;3. doi:10.3389/fphar.2012.00060

76. Hellstrom IC, Danik M, Luheshi GN, Williams S. Chronic LPS exposure produces changes in intrinsic membrane properties and a sustained IL-β-dependent increase in GABAergic inhibition in hippocampal CA1 pyramidal neurons. Hippocampus. 2005;15(5):656–64. doi:10.1002/hipo.20086

77. Hainmueller T, Bartos M. Dentate gyrus circuits for encoding, retrieval and discrimination of episodic memories. Nature Reviews Neuroscience. 2020 Mar 1;21(3):153–68. doi:10.1038/s41583-019-0260-z PubMed PMID: 32042144.

78. Hsu D. The dentate gyrus as a filter or gate: a look back and a look ahead. Prog Brain Res. 2007;163:601–13. doi:10.1016/S0079-6123(07)63032-5 PubMed PMID: 17765740.

79. Ewell LA. To sleep perchance to spike: a functional role for dentate spikes in memory. Journal of Physiology. 2017 Jul 15;595(14):4565. doi:10.1113/JP274502 PubMed PMID: 28485489.

80. Farrell JS, Hwaun E, Dudok B, Soltesz I. Neural and behavioural state switching during hippocampal dentate spikes [Internet]. doi:10.1038/s41586-024-07192-8

81. Flynn SP, Barrier S, Scott RC, Lenck-Santini PP, Holmes GL. Status epilepticus induced spontaneous dentate gyrus spikes: In vivo current source density analysis. PLoS ONE. 2015 Jul 6;10(7). doi:10.1371/journal.pone.0132630 PubMed PMID: 26148195.

82. Engel J, Bragin A, Staba R. Nonictal EEG biomarkers for diagnosis and treatment. Epilepsia Open. 2018 Dec 1;3(S2):120–6. doi:10.1002/epi4.12233

83. Perucca P, Smith G, Santana-Gomez C, Bragin A, Staba R. Electrophysiological biomarkers of epileptogenicity after traumatic brain injury. Neurobiology of Disease. 2019 Mar;123:69–74. doi:10.1016/j.nbd.2018.06.002

84. Tewari BP, Chaunsali L, Campbell SL, Patel DC, Goode AE, Sontheimer H. Perineuronal nets decrease membrane capacitance of peritumoral fast spiking interneurons in a model of epilepsy. Nature Communications. 2018 Dec 1;9(1). doi:10.1038/s41467-018-07113-0 PubMed PMID: 30413686.

85. Whitebirch AC, LaFrancois JJ, Jain S, Leary P, Santoro B, Siegelbaum SA, et al. Enhanced excitability of the hippocampal CA2 region and its contribution to seizure activity in a mouse model of temporal lobe epilepsy. Neuron. 2022 Oct;110(19):3121–3138.e8. doi:10.1016/j.neuron.2022.07.020

86. Zhou Y, Chen Y, Xu C, Zhang H, Lin C. TLR4 Targeting as a Promising Therapeutic Strategy for Alzheimer Disease Treatment. Front Neurosci. 2020 Dec 18;14:602508. doi:10.3389/fnins.2020.602508

87. Iori V, Iyer AM, Ravizza T, Beltrame L, Paracchini L, Marchini S, et al. Blockade of the IL-1R1/TLR4 pathway mediates disease-modification therapeutic effects in a model of acquired epilepsy. Neurobiology of Disease. 2017 Mar;99:12–23. doi:10.1016/j.nbd.2016.12.007

