## Supplementary Figure 1 for "Neuronal Toll-like Receptor-4 regulation of Matrix Metalloproteinase-9 Activity Mediates Dentate Circuit Dysfunction after Traumatic Brain Injury"

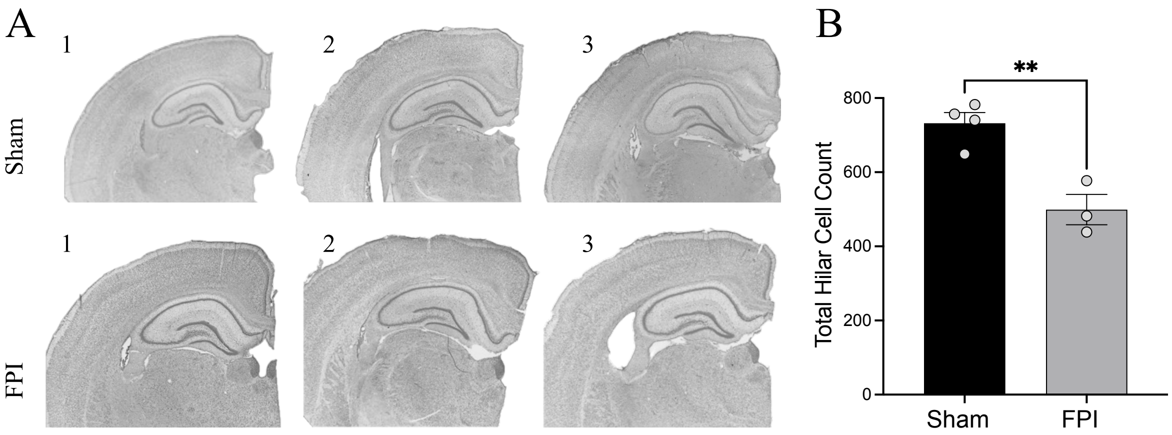


**Supplementary Figure 1: Absence of overt histopathological changes and consistent dentate gyrus hilar cell loss after Lateral Fluid Percussion Injury.** A) Representative images of Nissl-stained slices from 3 sham and 3 FPI rats 1-month post-injury showing no gross structural damage. B) Summary plot shows injury-induced decrease in hilar neuron count. n = 4 sham and 3 FPI rats. ** indicates P<0.01 by unpaired t-test.
