## Supplementary Figure 2 for "Neuronal Toll-like Receptor-4 regulation of Matrix Metalloproteinase-9 Activity Mediates Dentate Circuit Dysfunction after Traumatic Brain Injury"

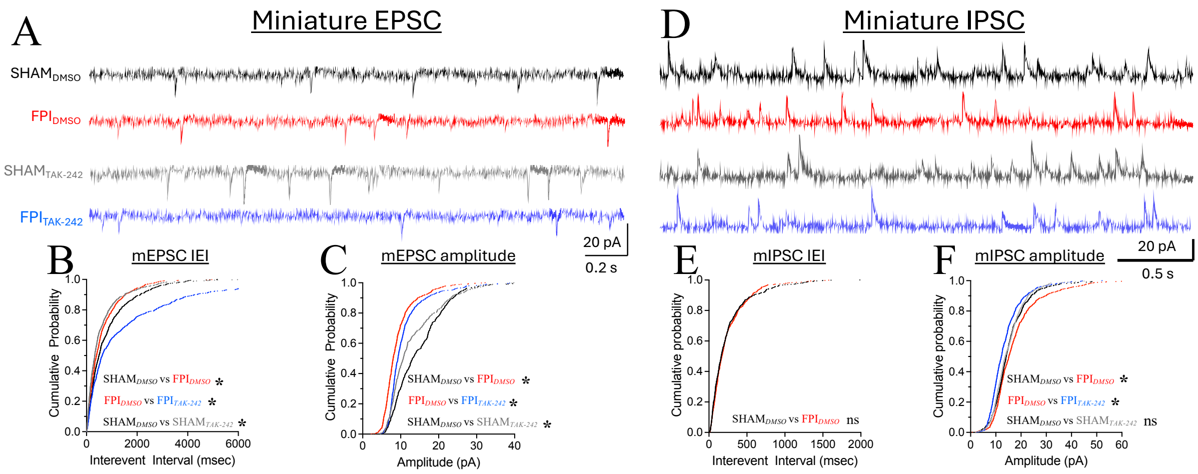


**Supplementary Figure 2: *TLR4 signaling drives changes in miniature excitatory and inhibitory inputs to DGCs after brain injury.*** A) Representative granule cell mEPSC traces from vehicle treated sham controls (black), vehicle treated FPI (red), TLR4 inhibited sham (grey) and TLR4 inhibited FPI (blue) rats. B-C) Cumulative probability plots of mEPSC interevent intervals (B) and amplitude (C) in experimental groups. D) Representative mIPSC traces from granule cells in vehicle treated sham controls (black), vehicle treated FPI (red), TLR4 inhibited sham (grey) and TLR4 inhibited FPI (blue) rats. E-F) Cumulative probability plots of mIPSC interevent intervals (E) and amplitude(F) in vehicle treated and TLR4 inhibited sham and injured rats. $n$ = 7 cells from 3 rats/group *indicates $P$<0.05 by Holm-Bonferroni corrected K.S. test.
