## Supplementary Figure 3 for "Neuronal Toll-like Receptor-4 regulation of Matrix Metalloproteinase-9 Activity Mediates Dentate Circuit Dysfunction after Traumatic Brain Injury"

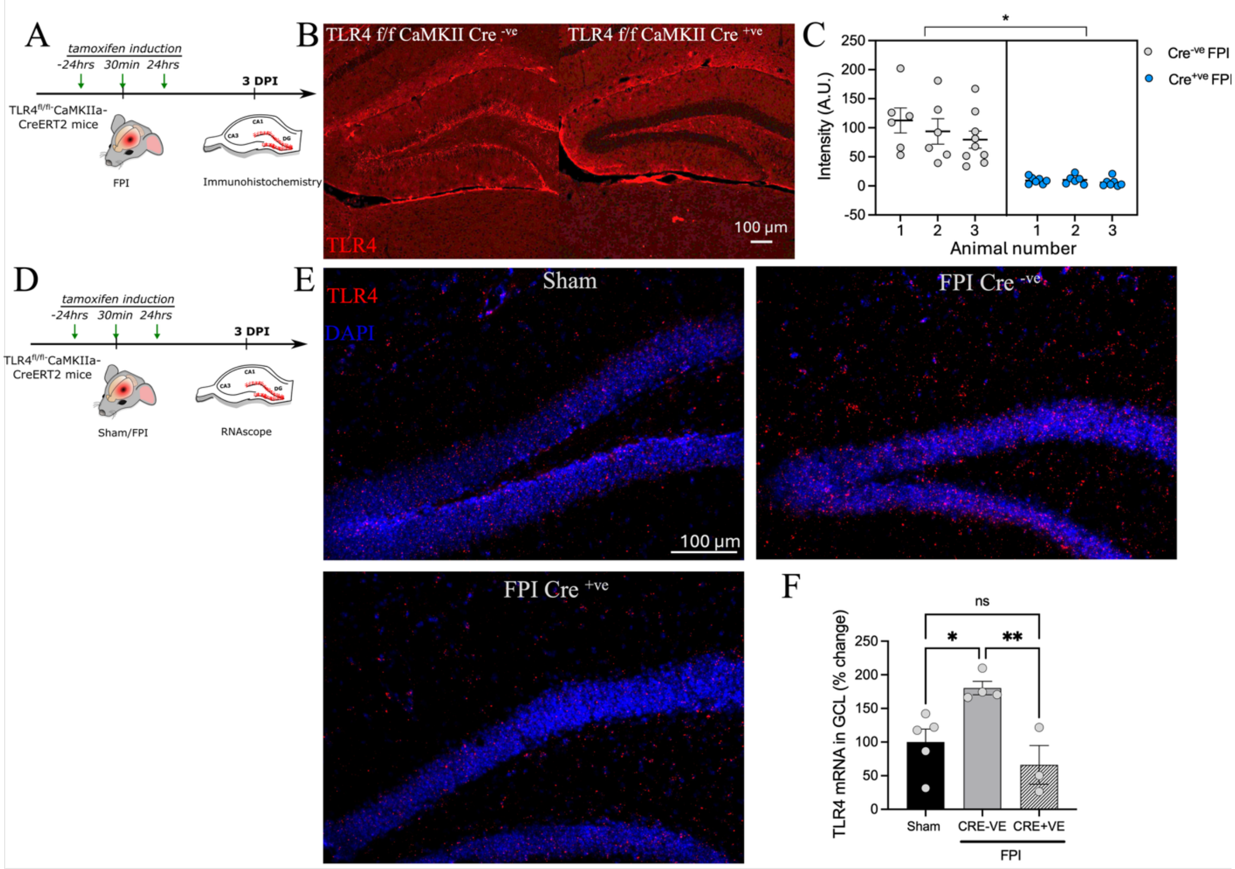


**Supplementary Figure 3:** ***Validation of cell-type specific TLR4 deletion in DG glutamatergic neurons.*** A) Experimental timeline for evaluating TLR4 expression after tamoxifen induction in TLR4$^{fl/fl}$: CaMKIIa-CreERT2 mice and littermate Cre$^{-ve}$ mice. B) Representative images of TLR4 expression in the DG of tamoxifen treated to Cre$^{-ve}$ and TLR4$^{fl/fl}$: CaMKIIa-CreERT2 mice 3 days after injury. C) Summary plot of TLR4 expression in experimental groups. * indicates P<0.05 by nested t-test based on n = 5-6 slices / mouse (indicated as animal number on x-axis) from 3 mice/group. D) Experimental timeline for evaluating TLR4 mRNA after tamoxifen induction in TLR4$^{fl/fl}$: CaMKIIa-CreERT2 mice and littermate Cre$^{-ve}$ mice. E) Representative images of TLR4 mRNA expression in the DG of a wild-type sham mouse, tamoxifen treated to Cre$^{-ve}$ and TLR4$^{fl/fl}$: CaMKIIa-CreERT2 mice 3 days after injury and a negative control. F) Summary plot of TLR4 mRNA expression in the granule cell layer (GCL) in experimental groups. * indicates $P<0.05$ and ** indicates $P<0.01$ by OW-ANOVA, based on data averaged across sections from n = 3-5 mice/group.
