## Supplementary Figure 4 for "Neuronal Toll-like Receptor-4 regulation of Matrix Metalloproteinase-9 Activity Mediates Dentate Circuit Dysfunction after Traumatic Brain Injury"

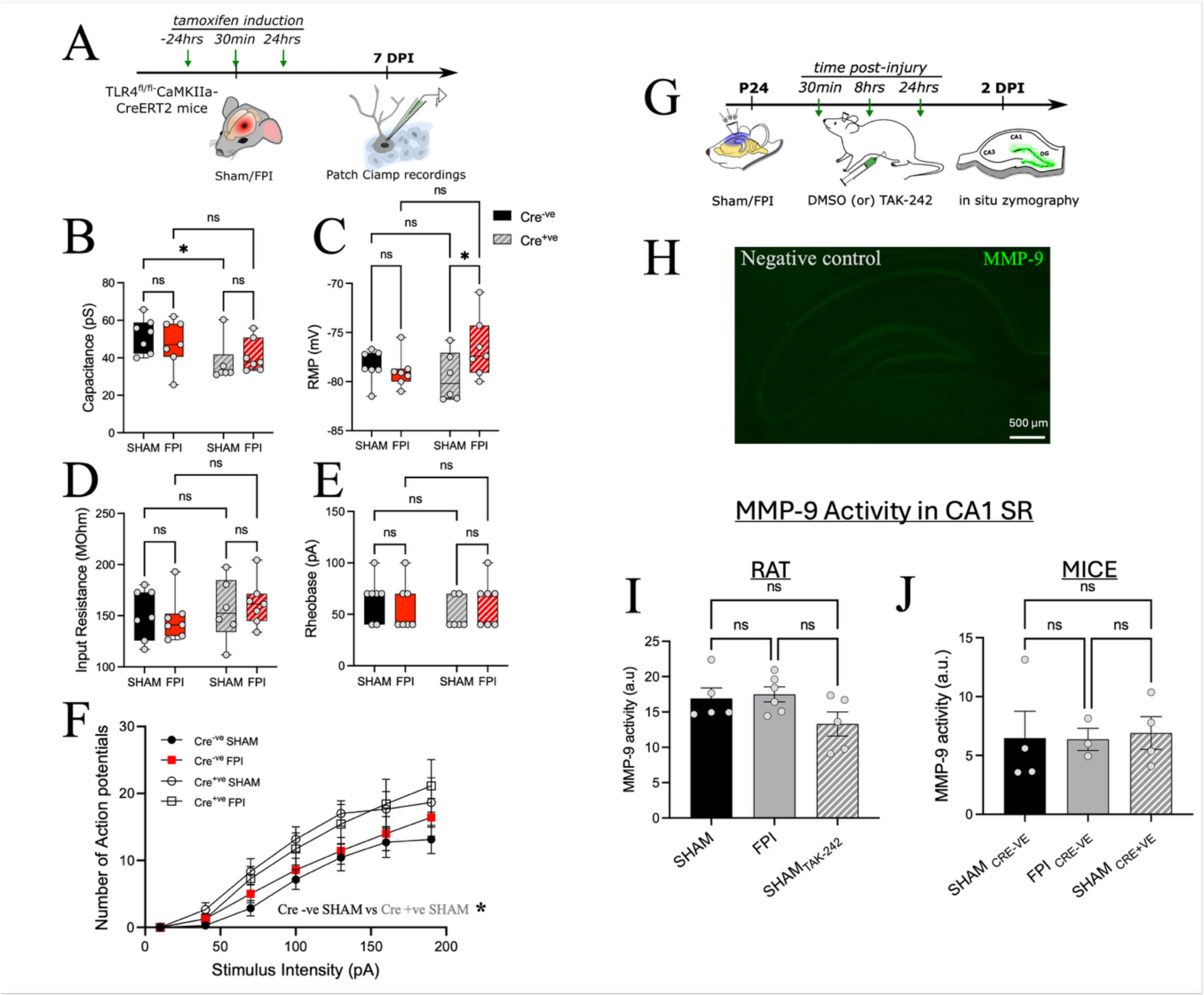


**Supplementary Figure 4:** ***Effects of selective TLR4 deletion in glutamatergic neurons on DGC properties.*** A) Experimental timeline for examining active and passive properties of DGCs from TLR4$^{fl/fl}$: CaMKIIa-CreERT2 mice. B-F) Summary plots of active and passive properties including capacitance (B), resting membrane potential (C), input resistance (D), Rheobase (E) and spike frequency (F) in Cre$^{-ve}$ and Cre$^{+ve}$ sham and FPI mice. Note the effect of genotype on cell capacitance and firing frequency in sham injured mice. Also note that the resting membrane potential is significantly depolarized Cre$^{+ve}$ mice after injury, but no differences are seen in Cre$^{-ve}$ mice. Data points in panels B-E represent individual cells (n = 7 cells from 3 mice/group). G)

Experimental timeline for *in situ* zymography experiments. H) Representative images from slices incubated in MMP-9 blocker SB-3CT that served as negative controls. I-J) Quantification of DQ-fluorescence showing CA1 Str. Radiatum intensity is unaltered by injury, treatments or genotype. * indicates $P<0.05$ by TW-ANOVA with Šídáks multiple comparisons post hoc tests (B-F) and OW-ANOVA (I-J). Data points in panel I and J represent individual animals (n = 5-6 rats/group, and 3-4 mice / group, respectively).
