## Supplementary Figure 5 for "Neuronal Toll-like Receptor-4 regulation of Matrix Metalloproteinase-9 Activity Mediates Dentate Circuit Dysfunction after Traumatic Brain Injury"

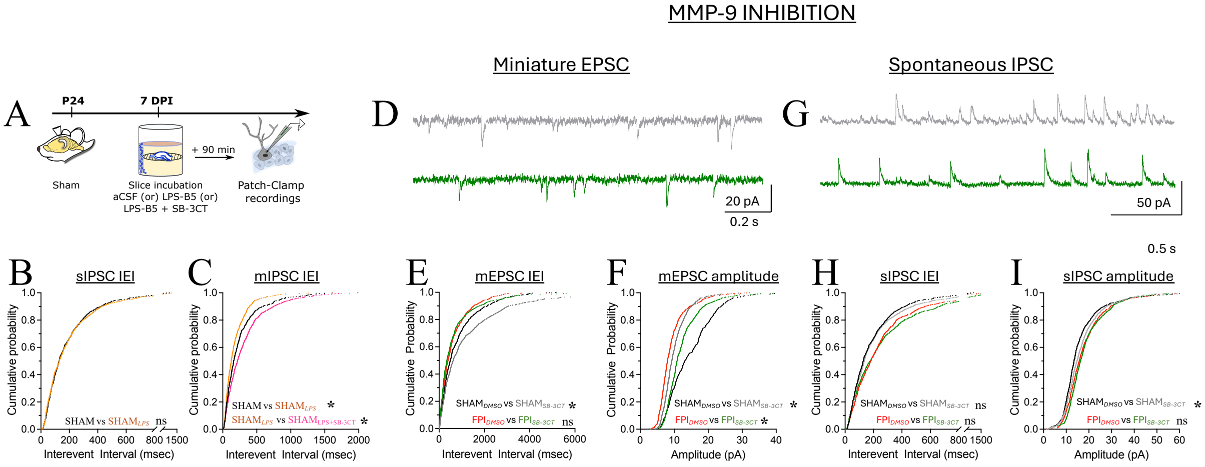


**Supplementary Figure 5:** ***MMP-9 inhibition modulates mEPSC amplitude without rescuing changes in sIPSC frequency or amplitude.*** A) Experimental timeline for examining changes in synaptic inputs to DGCs in acute rat slices after LPS incubation. B-C) Cumulative probability plots of granule cell sIPSC (B) and mIPSC (C) interevent intervals in sham, LPS-B5 incubated (orange) and LPS-B5+SB-3CT (magenta) co-incubated slices. Note that while LPS-B5 incubation did not alter sIPSC, mIPSC interevent intervals were significantly reduced. Co-incubation of slices in MMP-9 blocker SB-3CT prevented this change. D) Representative granule cell mEPSC traces from SB-3CT treated sham (grey) and injured rats (green). E-F) Cumulative probability plots of mEPSC interevent intervals (E) and amplitude (F) in SB-3CT treated sham/injured rats. G) Representative granule cell sIPSC recordings from SB-3CT treated sham (grey) and injured rats (green). H-I) Cumulative probability plots of mEPSC interevent intervals (H) and amplitude (I) between SB-3CT treated sham/injured rats. n = 8 cells from 4 rats/group (A-C), 7-9 cells from 3 rats/group (D-F) and 8-13 cells from 3 rats/group (G-I) *indicates $P$<0.05 by Holm-Bonferroni corrected K.S. test.
