## Supplementary Figure 6 for "Neuronal Toll-like Receptor-4 regulation of Matrix Metalloproteinase-9 Activity Mediates Dentate Circuit Dysfunction after Traumatic Brain Injury"

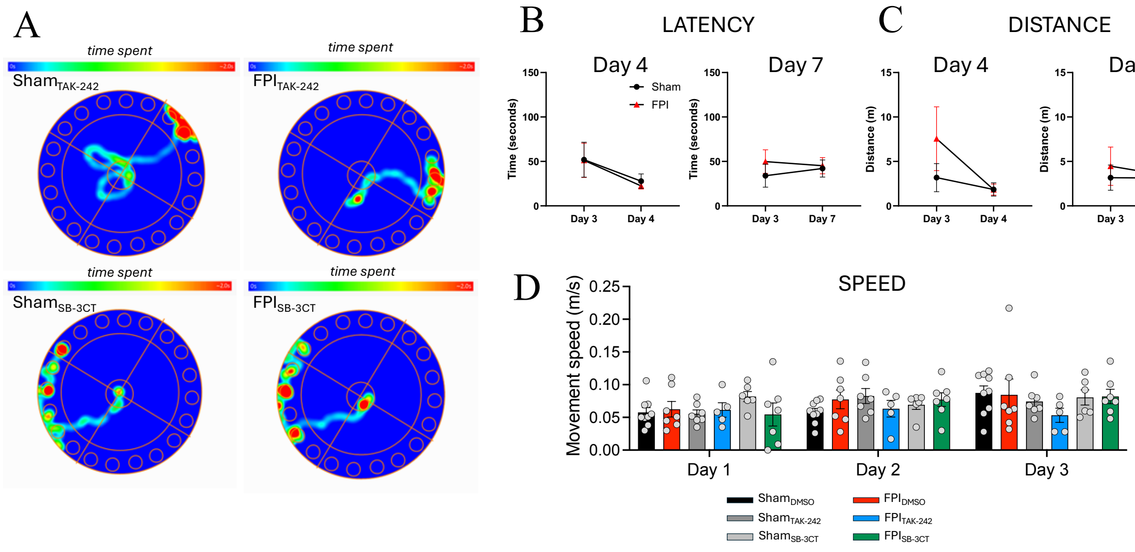


**Supplementary Figure 6:** ***Treatment with Tak-242 or SB-3CT mitigates Spatial learning deficits in brain injured rats.*** A) Representative heat plots showing trajectory of drug-treated rats in the Barnes maze during task on day 2. B-C) Summary plot of latency to escape (B) and distance travelled (C) on Barnes maze spatial navigation task on day 4 or day 7 in untreated sham and injured rats. D) Summary plot of speed of movement across the training days between the vehicle and drug-treated sham and injured animals showed no injury or training day differences. n = 6-10 rats/group. Data points in panel D represent individual animals.
