## Supplementary Table1 for "Neuronal Toll-like Receptor-4 regulation of Matrix Metalloproteinase-9 Activity Mediates Dentate Circuit Dysfunction after Traumatic Brain Injury"

TABLE 1:

| *Spontaneous EPSC* | | | | | | | |
|---|---|---|---|---|---|---|---|
| | **Interevent intervals** | | | **Amplitude** | | | |
| | **Mean (ms)** | **Median** | **Interquartile range** | **Mean (pA)** | **Median** | **Interquartile range** | ***n* (cells)** |
| $SHAM_{DMSO}$ | 336.9 | 216.4 | 86.05 - 429.9 | 13.16 | 12.20 | 9.99-15.36 | 8 |
| $FPI_{DMSO}$ | 225.3 | 144.8 | 64.60 - 307.4 | 11.23 | 10.31 | 8.06-13.42 | 8 |
| $SHAM_{TAK-242}$ | 221.8 | 143.6 | 65.74 - 285.6 | 13.20 | 12.23 | 9.64-15.77 | 9 |
| $FPI_{TAK-242}$ | 321.9 | 204.2 | 95.08 - 420.2 | 13.82 | 12.49 | 9.80-16.20 | 8 |
| $SHAM_{LPS-B5}$ | 265.3 | 157.3 | 69.18 - 330.4 | 17.86 | 17.29 | 14.06-21.09 | 8 |
| $SHAM_{LPS-B5+SB-3CT}$ | 339.7 | 230.9 | 94.53 - 434.6 | 15.90 | 14.66 | 11.53-18.96 | 9 |
| $SHAM_{SB-3CT}$ | 311.3 | 203.5 | 81.18 - 396.8 | 12.45 | 11.38 | 9.22-14.57 | 10 |
| $FPI_{SB-3CT}$ | 495.6 | 303.8 | 119.8 - 632.3 | 12.60 | 11.64 | 9.29-14.88 | 9 |
| *Spontaneous IPSC* | | | | | | | |
| $SHAM_{DMSO}$ | 190.3 | 128.1 | 65.35 - 242.4 | 16.66 | 13.78 | 10.81-19.21 | 8 |
| $FPI_{DMSO}$ | 256.6 | 181.1 | 83.78 - 320.4 | 18.24 | 15.74 | 11.94-21.70 | 8 |
| $SHAM_{TAK-242}$ | 174.2 | 102.4 | 50.75 - 199.2 | 14.79 | 12.98 | 10.27-16.85 | 8 |
| $FPI_{TAK-242}$ | 200.9 | 119.7 | 61.08 - 252 | 19.95 | 14.18 | 10.58-23.23 | 7 |
| $SHAM_{LPS-B5}$ | 197.8 | 128.6 | 63.33 - 247.5 | 15.56 | 13.52 | 10.99-17.58 | 8 |
| $SHAM_{LPS-B5+SB-3CT}$ | 294.3 | 206.4 | 92.83 - 395.6 | 18.01 | 16.17 | 12.30-21.37 | 10 |
| $SHAM_{SB-3CT}$ | 208.3 | 135.9 | 69.60 - 255.9 | 16.96 | 14.77 | 11.16-20.12 | 13 |
| $FPI_{SB-3CT}$ | 288.3 | 177.2 | 73.75 - 361.9 | 18.71 | 16.28 | 12.78-21.90 | 8 |
| **TLR4fl/fl-CaMKIIa-CreERT2 mice – Spontaneous EPSC** | | | | | | | |
| | **Mean (sec)** | **Median** | **Interquartile range** | **Mean (pA)** | **Median** | **Interquartile range** | ***n* (cells)** |
| $Cre^{-ve}$ SHAM | 2.154 | 1.291 | 0.45-2.74 | 13.42 | 11.65 | 10.03-14 | 7 |
| $Cre^{-ve}$ FPI | 1.234 | 0.721 | 0.31-1.79 | 11.29 | 10.36 | 8.62-12.73 | 7 |
| $Cre^{-ve}$ $FPI_{SB-3CT}$ | 1.734 | 0.997 | 0.36-2.24 | 10.05 | 9.222 | 7.57-11.43 | 7 |
| $Cre^{+ve}$ SHAM | 2.223 | 1.271 | 0.49-3.10 | 12.01 | 11.19 | 8.90-13.62 | 6 |
| $Cre^{+ve}$ FPI | 1.904 | 1.203 | 0.41-2.46 | 11.27 | 10.42 | 8.46-12.65 | 7 |

**Table 1. Descriptive statistics dentate gyrus cells (DGCs) physiology following traumatic brain injury.** Summary of descriptive statistics of interevent intervals and amplitude of spontaneous excitatory and inhibitory postsynaptic currents recorded from DGCs. n represents cells recorded from 3-4 animals/group.
