## Supplementary Table2 for "Neuronal Toll-like Receptor-4 regulation of Matrix Metalloproteinase-9 Activity Mediates Dentate Circuit Dysfunction after Traumatic Brain Injury"

TABLE 2:

| *Miniature EPSC* | | | | | | | |
|---|---|---|---|---|---|---|---|
| | **Interevent intervals** | | | **Amplitude** | | | |
| | **Mean (ms)** | **Median** | **Interquartile range** | **Mean (pA)** | **Median** | **Interquartile range** | ***n* (cells)** |
| $\text{SHAM}_{\text{DMSO}}$ | 824.8 | 496.1 | 208.1 - 1096 | 14.16 | 12.32 | 8.90-17.91 | 7 |
| $\text{FPI}_{\text{DMSO}}$ | 642.4 | 396.4 | 158.2 - 858.6 | 9.28 | 8.02 | 6.55-10.55 | 7 |
| $\text{SHAM}_{\text{TAK-242}}$ | 615.2 | 325.4 | 130 - 749.2 | 12.24 | 10.54 | 8.15-15.13 | 8 |
| $\text{FPI}_{\text{TAK-242}}$ | 1648 | 621.2 | 234.5 - 1959 | 13.82 | 12.49 | 9.80-16.20 | 8 |
| $\text{SHAM}_{\text{LPS-B5}}$ | 726.5 | 467.7 | 189.6 - 982.4 | 10.04 | 9.219 | 7.37-11.74 | 6 |
| $\text{SHAM}_{\text{LPS-B5+SB-3CT}}$ | 1014 | 551.1 | 197.5 - 1272 | 12.75 | 11.23 | 9.53-14.44 | 10 |
| $\text{SHAM}_{\text{SB-3CT}}$ | 1313 | 624.1 | 243.6 - 1660 | 10.22 | 9.30 | 7.63-11.82 | 9 |
| $\text{FPI}_{\text{SB-3CT}}$ | 710.5 | 346.9 | 146.9 - 784.7 | 12.36 | 10.85 | 8.74-14.04 | 7 |
| *Miniature IPSC* | | | | | | | |
| | **Interevent intervals** | | | **Amplitude** | | | |
| | **Mean (ms)** | **Median** | **Interquartile range** | **Mean (pA)** | **Median** | **Interquartile range** | ***n* (cells)** |
| $\text{SHAM}_{\text{DMSO}}$ | 254.7 | 161.2 | 77.05 - 318.9 | 15.72 | 14.15 | 10.99-18.58 | 7 |
| $\text{FPI}_{\text{DMSO}}$ | 251 | 171 | 86.30 - 336.7 | 17.68 | 15.20 | 11.58-20.72 | 7 |
| $\text{SHAM}_{\text{TAK-242}}$ | 217.6 | 150.1 | 74.63 - 286.1 | 15.14 | 13.72 | 10.54-18.27 | 7 |
| $\text{FPI}_{\text{TAK-242}}$ | 251 | 143 | 64.70 - 300.6 | 13.75 | 12.23 | 9.26-16.35 | 6 |
| $\text{SHAM}_{\text{LPS-B5}}$ | 183.8 | 124.9 | 60.13 - 244.1 | 14.06 | 12.56 | 10.04-16.53 | 6 |
| $\text{SHAM}_{\text{LPS-B5+SB-3CT}}$ | 321.6 | 216.9 | 90.60 - 420 | 18.05 | 16.33 | 13.19-21.49 | 10 |
| $\text{SHAM}_{\text{SB-3CT}}$ | 218.1 | 140 | 63.63 - 287.2 | 15.95 | 14.31 | 11.41-18.37 | 11 |
| $\text{FPI}_{\text{SB-3CT}}$ | 366.4 | 208.1 | 93.60 - 462.2 | 16.67 | 14.90 | 11.47-19.80 | 7 |

**Table 2. Descriptive statistics dentate gyrus cells (DGCs) physiology following traumatic brain injury.** Summary of descriptive statistics of interevent intervals and amplitude of miniature excitatory and inhibitory postsynaptic currents recorded from DGCs. n represents cells recorded from 3-4 animals/group.
