## Supplementary Table3 for "Neuronal Toll-like Receptor-4 regulation of Matrix Metalloproteinase-9 Activity Mediates Dentate Circuit Dysfunction after Traumatic Brain Injury"

TABLE 3: *Holm-Šídák Adjusted P-values*

| | **Excitatory currents** | | | |
|---|---|---|---|---|
| | **Spontaneous** | | **Miniature** | |
| | IEI | amplitude | IEI | amplitude |
| $SHAM_{DMSO}$ (vs) $FPI_{DMSO}$ | *P*<0.0001; d = 0.344 | *P*<0.0001; d = 0.410 | *P*=0.0027; d =0.217 | *P*<0.0001; d = 0.853 |
| $FPI_{DMSO}$ (vs) $FPI_{TAK\text{-}242}$ | *P*<0.0001; d = 0.615 | *P*<0.0001; d =0.502 | *P*<0.0001; d =0.543 | *P*<0.0001; d = 0.889 |
| $SHAM_{DMSO}$ (vs) $SHAM_{TAK\text{-}242}$ | *P*<0.0001; d = 0.354 | *P=0.5297*; d = 0.008 | *P*<0.0001; d =0.239 | *P*<0.0001; d = 0.301 |
| $SHAM_{DMSO}$ (vs) $Sham_{LPS}$ | *P*<0.0001; d = 0.202 | *P*<0.0001; d = 0.766 | *P*=0.2463; d = 0.113 | *P*<0.0001; d = 0.788 |
| $Sham_{LPS}$ (vs) $Sham_{LPS + SB\text{-}3CT}$ | *P*<0.0001; d = 0.216 | *P*<0.0001; d = 0.325 | *P*=0.0012; d = 0.237 | *P*<0.0001; d = 0.538 |
| $SHAM_{DMSO}$ (vs) $Sham_{SB\text{-}3CT}$ | *P* =0.6476; d = 0.069 | *P* =0.0002; d = 0.147 | *P* =0.0001; d = 0.311 | *P*<0.0001; d = 0.738 |
| $FPI_{DMSO}$ (vs) $FPI_{SB\text{-}3CT}$ | *P*<0.0001; d = 0.601 | *P*<0.0001; d = 0.301 | *P* =0.1108; d = 0.073 | *P*<0.0001; d = 0.620 |
| | **Inhibitory currents** | | | |
| $SHAM_{DMSO}$ (vs) $FPI_{DMSO}$ | *P*<0.0001; d = 0.290 | *P*<0.0001; d = 0.157 | *P*=0.5139; d = 0.013 | *P*=0.0044; d = 0.220 |
| $FPI_{DMSO}$ (vs) $FPI_{TAK\text{-}242}$ | *P*<0.0001; d = 0.221 | *P*<0.0001; d = 0.112 | *P*=0.0748; d =0.000 | *P*<0.0001; d = 0.473 |
| $SHAM_{DMSO}$ (vs) $SHAM_{TAK\text{-}242}$ | *P*=0.0005; d = 0.072 | *P*=0.0037; d = 0.206 | *P*=0.5139; d = 0.419 | *P*=0.3 808; d = 0.077 |
| $SHAM_{DMSO}$ (vs) $Sham_{LPS}$ | *P*=0.8642; d = 0.037 | *P*=0.0222; d = 0.107 | *P*=0.0004; d = 0.293 | *P*<0.0001; d =0.226 |
| $Sham_{LPS}$ (vs) $Sham_{LPS + SB\text{-}3CT}$ | *P*<0.0001; d =0.378 | *P*<0.0001; d = 0.246 | *P*<0.0001; d = 0.468 | *P*<0.0001; d = 0.549 |
| $SHAM_{DMSO}$ (vs) $Sham_{SB\text{-}3CT}$ | *P*=0.4758; d = 0.085 | *P*<0.0001; d =0.138 | *P*=0.0542; d = 0.146 | *P*=0.2826; d =0.030 |
| $FPI_{DMSO}$ (vs) $FPI_{SB\text{-}3CT}$ | *P*=0.4287; d = 0.098 | *P*=0.0597; d = 0.048 | *P*=0.0012; d = 0.299 | *P*=0.5412; d = 0.113 |
| **TLR4fl/fl-CaMKIIa-CreERT2 mice – Spontaneous Excitatory currents** | | | | |
| $Cre^{-ve}$ SHAM (vs) $Cre^{-ve}$ FPI | *P*<0.0001; d = 0.420 | *P*<0.0001; d = 0.311 | | |
| $Cre^{+ve}$ SHAM (vs) $Cre^{+ve}$ FPI | *P*=0.1109; d = 0.131 | *P*=0.0008; d = 0.157 | | |
| $Cre^{-ve}$ FPI vs $Cre^{-ve}$ $FPI_{SB\text{-}3CT}$ | *P*<0.0001; d = 0.283 | *P*<0.0001; d = 0.303 | | |

**Table 3. Statistical analysis of synaptic inputs to dentate gyrus cells (DGCs) following traumatic brain injury.** Summary of Holm-Bonferroni corrected *P*-values (Kolmogorov-Smirnov tests) and Cohen's *d* effect sizes for comparisons of synaptic inputs to DGCs between sham and FPI groups. Animals received either vehicle or the TLR4 inhibitor TAK-242. Parameters include interevent intervals and amplitudes for both spontaneous and miniature excitatory postsynaptic currents (sEPSC/mEPSC) and inhibitory postsynaptic currents (sIPSC/mIPSC).
