## Supplementary Table4 for "Neuronal Toll-like Receptor-4 regulation of Matrix Metalloproteinase-9 Activity Mediates Dentate Circuit Dysfunction after Traumatic Brain Injury"

| | **10min** | **60min** | **120min** | **180min** |
|---|---|---|---|---|
| $SHAM_{DMSO}$ | 157.71±6.58 | 144.00±4.30 | 146.78±4.82 | 145.15±7.03 |
| $FPI_{DMSO}$ | 130.23±6.92 | 116.69±3.84 | 112.84±4.20 | 110.36±5.31 |
| $SHAM_{TAK\text{-}242}$ | 150.12±7.95 | 133.00±5.58 | 124.33±8.60 | 116.11±6.98 |
| $FPI_{TAK\text{-}242}$ | 149.14±8.95 | 143.91±4.66 | 140.32±4.02 | 141.21±5.00 |
| $SHAM_{SB\text{-}3CT}$ | 164.56±5.67 | 159.66±2.40 | 156.98±8.57 | 154.61±7.70 |
| $FPI_{SB\text{-}3CT}$ | 166.98±10.3 | 157.60±10 | 146.85±6.61 | 134.76±8.62 |

**Table 4. Maintenance of long-term potentiation (LTP) in the hippocampal dentate gyrus post-injury.** Normalized slope values (mean ± SEM) measured at 10, 60, 120, and 180 min following theta burst stimulation (TBS). Measurements were recorded one-week post-injury in sham and FPI rats treated with vehicle (DMSO), a TLR4 inhibitor (TAK-242), or an MMP-9 inhibitor (SB-3CT).
