## Supplementary Table5 for "Neuronal Toll-like Receptor-4 regulation of Matrix Metalloproteinase-9 Activity Mediates Dentate Circuit Dysfunction after Traumatic Brain Injury"

TABLE 5:

| | | **Day1** | | **Day2** | | **Day3** | |
|---|---|---|---|---|---|---|---|
| **Group** | *n* (rats) | Latency (seconds) | Distance (meters) | Latency (seconds) | Distance (meters) | Latency (seconds) | Distance (meters) |
| $SHAM_{DMSO}$ | 10 | 61.36±11.11 | 3.32±0.87 | 49.19±9.30 | 2.83±0.36 | 41.18±10.78 | 3.17±1.00 |
| $FPI_{DMSO}$ | 10 | 113.1±16.68 | 7.20±1.67 | 87.06±10.64 | 9.07±1.72 | 50.21±10.34 | 5.39±1.80 |
| $SHAM_{TAK\text{-}242}$ | 7 | 69.30±19.92 | 3.14±0.58 | 82.90±25.52 | 6.06±1.41 | 67.74±21.15 | 4.84±1.01 |
| $FPI_{TAK\text{-}242}$ | 7 | 102.2±24.2 | 5.25±1.10 | 40.26±11.01 | 2.76±0.46 | 58.04±16.63 | 3.53±1.18 |
| $SHAM_{SB\text{-}3CT}$ | 6 | 82.58±24.05 | 10.4±2.30 | 42.12±10.34 | 3.00±0.84 | 62.50±6.84 | 4.68±0.89 |
| $FPI_{SB\text{-}3CT}$ | 7 | 67.86±18.33 | 5.43±2.67 | 46.66±4.97 | 3.46±0.40 | 51.73±22.18 | 4.51±2.29 |

**Table 5. Spatial memory performance in the Barnes maze task one month after brain injury.** Mean latency (seconds) to reach the escape hole and total distance traveled (meters) by sham and FPI rats on training days 1–3. Testing was conducted one-month post-injury and values are given as mean ± SEM. Treatment groups received vehicle (DMSO), TLR4 inhibitor (TAK-242), or MMP-9 inhibitor (SB-3CT) immediately following injury.
